# CD4 T cell dysfunction is associated with bacterial recrudescence during chronic tuberculosis

**DOI:** 10.1101/2025.01.22.634376

**Authors:** Evelyn Chang, Kelly Cavallo, Samuel M. Behar

## Abstract

While most people contain *Mycobacterium tuberculosis* infection, some individuals develop active disease, usually within two years of infection. Why immunity fails after initially controlling infection is unknown. C57BL/6 mice control *Mycobacterium tuberculosis* for up to a year but ultimately succumb to disease. We hypothesize that the development of CD4 T cell dysfunction permits bacterial recrudescence. We developed a reductionist model to assess antigen-specific T cells during chronic infection and found evidence of CD4 T cell senescence and exhaustion. In C57BL/6 mice, CD4 T cells upregulate coinhibitory receptors and lose effector cytokine production. Single cell RNAseq shows that only a small number of CD4 T cells in the lungs of chronically infected mice are polyfunctional. While the origin and causal relationship between T-cell dysfunction and recrudescence remains uncertain, we propose T cell dysfunction leads to a feed-forward loop that causes increased bacillary numbers, greater T cell dysfunction, and progressive disease.

## Introduction

Despite the development of vaccines (1920’s), diagnostic tests (1930’s), and antibiotics (1940’s), tuberculosis (TB) remains one of the leading causes of death from a pathogen. Approximately 25% of the world’s population has been exposed to *Mycobacterium tuberculosis* (Mtb), the causative agent of TB, and of those, 5-10% develop disease ^1, 2^. In most cases, why individuals progress to active pulmonary disease is poorly understood ^3^. However, data from HIV-infected people and animal models consistently find that CD4 T cells are crucial for immunity against Mtb ^4–6^. Given the importance of CD4 T cells in controlling Mtb infection and the lack of knowledge about how infection and its accompanying chronic inflammation affect the function of CD4 T cells, we hypothesize that bacterial recrudescence and the development of active disease, which can occur in some people after long periods of effective immune-mediated control, is the consequence of CD4 T cell dysfunction.

T cell exhaustion occurs when persistent antigen presentation leads to chronic T cell stimulation. Chronically activated T cells undergo transcriptional changes, including increased co-inhibitory receptor (i.e. PD-1, CTLA-4, TOX, TIM-3, LAG-3, TIGIT) expression and decreased effector function ^7^. Currently, CD8 T cells in cancer or chronic viral infections are the focus of most studies of T cell exhaustion, and these studies have led to the development of monoclonal antibody (mAb) blockade treatments to reinvigorate T cells in the tumor microenvironment ^8^. More focused studies revealed distinct subpopulations exhausted T-cells: progenitor exhausted T cells with stem-like properties that respond to mAb blockade and terminally exhausted T cells that are non-proliferative and unresponsive to mAb treatment ^9–13^. The relative abundance of these populations within the tumor microenvironment can inform disease prognosis and guide therapy ^14, 15^.

While CD4 T cell exhaustion has been described ^7, 8^, it is one of several types of dysfunction that has been identified during CD4 T cell responses. Other CD4 dysfunctional states include anergy, senescence, and activation-induced cell death ^16, 17^. Compared to CD8 T cells, much less is known about how CD4 T cells respond to chronic TCR stimulation. It is unclear whether current definitions of CD8 T cell exhaustion apply to CD4 T cells, especially during chronic bacterial infections. As TB is a chronic bacterial infection, and the key protective immune responses are mediated by CD4 T cells, the murine TB model provides a unique opportunity to study CD4 T cell exhaustion during chronic bacterial infection. Using an adoptive transfer model, we established that CD4 T cells in the chronically infected lung have an exhausted phenotype based on cell surface expression of coinhibitory receptors, loss of function, and a transcriptomic profile resembling exhausted CD8 T cells. These features were associated with an increased bacterial burden and premature death. In chronically infected C57BL/6 (B6) mice, parenchymal CD4 T cells express PD-1 and TIM-3 coinhibitory receptors and have a diminished capacity to produce cytokines; thus, they resemble terminally exhausted CD8 T cells. Single cell RNAseq analysis revealed that many CD4 clusters expressed *Ifng*, only two clusters appeared appeared to be truly polyfunctional with respect to cytokine production. Late during chronic infection, a single small cluster of CD4 T cells remained polyfunctional. Senescence was identified as a cellular phenotype that contributes to CD4 dysfunction in both the adoptive transfer model and in B6 mice. Hypoxia and TCR affinity appear to modulate CD4 T cell dysfunction. Further study of the factors and mechanisms that promote T cell dysfunction could inform the development of therapeutic options to protect T cells and maintain their function.

## Results

### A reductionist model identifies CD4 T cell dysfunction during chronic Mtb infection

To study antigen-specific CD4 T cells in the setting of chronic Mtb infection, we developed an adoptive transfer model (Figure 1a). We observed that intravenous transfer of P25 (i.e., Ag85b-specific CD4) TCR transgenic (TCRtg) T cells to TCRα ko mice prolonged the median survival time (MST) of recipient mice from five weeks to 18 weeks (Figure 1b) ^18^. Under the same infection conditions, B6 mice survive >48 weeks ^19, 20^. P25 T cells gradually lose the capacity for cytokine production during chronic infection, but the diminished expression of Ag85b by Mtb in vivo raised the possibility that recrudescence was secondary to the inability of P25 T cells to recognize Mtb-infected cells ^21^. Therefore, we transferred C7 (i.e., ESAT-6- specific CD4) TCRtg T cells into TCRα KO mice (C7➔TCRα). C7 T cells conferred long-term protection, but ultimately the mice succumbed with an MST of 26 weeks, which is significantly reduced compared to C57BL/6 mice (Figure 1b). The greater susceptibility of C7➔TCRα mice was accompanied by a progressive increase in lung bacterial burden (Figure 1c). We used this model to investigate why immunity failed.

**Figure 1.**
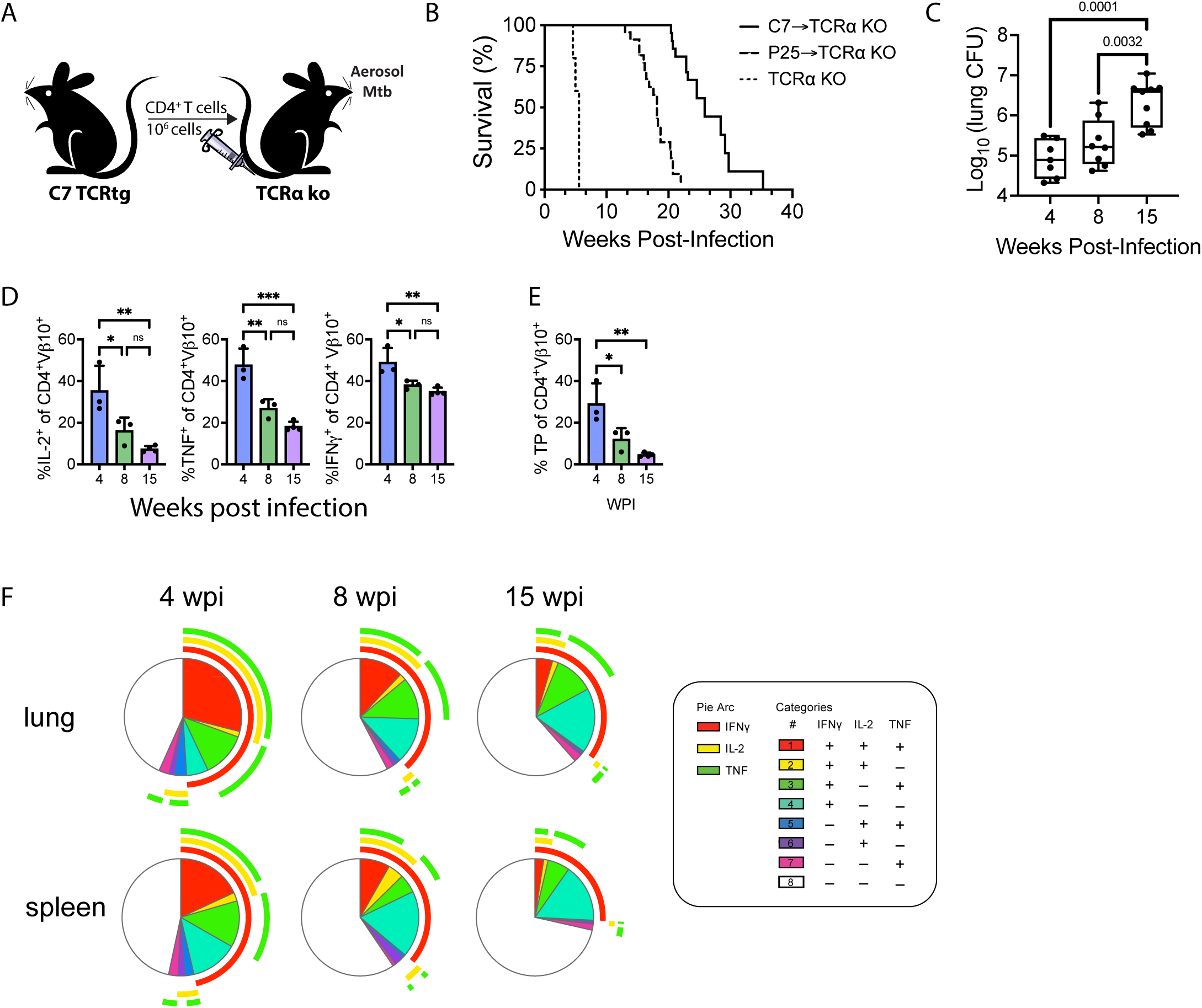
Loss of CD4 T cell function during chronic Mtb infection is associated with recrudescence and death in the C7➔TCRα adoptive transfer model. (A) Schematic of C7➔TCRα adoptive transfer model. (B) Survival curves of infected TCRα KO, P25➔TCRα KO (n=20), and C7➔TCRα KO (n=13). The difference between the groups was statistically significant (p < 0.0001) as determined by the log-rank test. (C) Lung CFU of infected C7➔TCRα mice at 4, 8, and 15 wpi. Data is pooled from two independent experiments, each with 3-5 mice per time point. Box plots indicate median (middle line), 25th, 75th percentile (box) and minimum and maximum (whiskers). (D) Frequencies of IL-2, TNF, or IFNγ or (E) Triple Producers (TP; IL- 2^+^TNF^+^IFNγ^+^) producing C7 T cells. Representative data of three independent experiments, 3-5 mice/group. Bars, mean ± SD. (F) Cytokine production and T cell polyfunctionality visualized using SPICE plots. (C-E) Statistical significance was analyzed by one-way ANOVA with Bonferroni’s multiple comparison test. p-values: *, p<0.05; **, p<0.01; ***, p<0.001; ****, p<0.0001; ns, no significant difference.

An advantage of the C7➔TCRα model is the large number of antigen-specific T cells that can be assessed independently of their function. Purified CD4 T cells from C7 mice contained variable numbers of non-transgenic naïve CD4 T cells (see methods), so ESAT6- specific CD4 T cells were identified using ESAT6/IA^b^ (E6) tetramers or anti-Vβ10 antibodies (see Figure S1a). Cytokine production in response to ESAT6 peptide stimulation was used as a measure of T cell function. The frequency of C7 T cells producing IL-2, TNF, or IFNγ decreased over time, accompanied by a loss of polyfunctional T cells and an increase of non-cytokine-producing T cells (Figure 1d, 1e, 1f). Thus, diminished function of ESAT6-specific CD4 T cells correlates with increasing bacillary burden.

To determine whether loss of function was associated with the acquisition of co-inhibitory receptor expression, PD-1, TIM-3, LAG-3, CTLA-4, and 2B4 expression was measured between 4-and 15-weeks post-infection (wpi). PD-1 and TIM-3 were the most highly expressed, and the number of co-inhibitory receptors expressed by C7 T cells increased over time (Figure 2a, 2b). The frequency of PD-1^+^TIM-3^+^ C7 T cells increased over time, a characteristic of terminally exhausted CD8 T cells (Figure 2c) ^13, 22^. In parallel, C7 T cells resembling progenitor exhausted CD8 T cells (T_PEX_; i.e., TIM-3^-^TCF-1^+^ or TIM-3^-^SLAMF6^+^) diminished while those resembling terminally exhausted CD8 T cells (T_TEX_; i.e., TIM-3^+^TCF-1^-^ or TIM-3^+^SLAMF6^-^) increased (Figure 2d, 2e) ^13, 22^. The percentage of TIM-3^+^TCF-1^-^ C7 T cells expressing PD-1 was higher compared to their TIM-3^-^TCF-1^+^ counterpart (Figure 2f). Consistent with a terminally exhausted phenotype, PD-1 levels were significantly higher on TIM-3^+^TCF-1^-^ C7 T cells (Figure 2g). Although the kinetics of the expression of co-inhibitory receptors varied between experiments, we always observed CD4 T cells expressing high levels of PD-1 and TIM-3 late during infection We next asked whether there was a correlation between the exhausted phenotype of C7 cells and progressive lung infection. Analysis of all individuals from two independent experiments encompassing multiple time points demonstrated a strong positive correlation between the frequency of PD-1^+^ C7 T cells and CFU (Figure 2h, left). As PD-1 is also a marker of activated T cells, we next determined the correlation between PD-1^+^TIM-3^-^ (i.e., activated or T_PEX_) and PD-1^+^TIM-3^+^ (T_TEX_) C7 cells. There was no significant correlation between the PD-1^+^TIM-3^-^ phenotype and lung CFU; in contrast, there was a striking correlation with the PD-1^+^TIM-3^+^ C7 phenotype, which was largely independent of time point or experiment (Figure 2h). While the high number of C7 cells could create competition for antigen and APC in this model, we suggest that this would have the opposite effect on T cell exhaustion: namely, that less TCR stimulation would preserve function. These data suggest that the emergence of CD4 T cell exhaustion late during infection could contribute to recrudescence and disease. However, the causal relationship remains to be established. Development of T cell exhaustion could result in loss of Mtb control, or alternatively, an increase in bacillary burden and antigen load could result in T cell exhaustion. Nevertheless, these data show that the development of terminally exhausted CD4 T cells correlates with ongoing bacterial replication and disease progression in this model.

**Figure 2.**
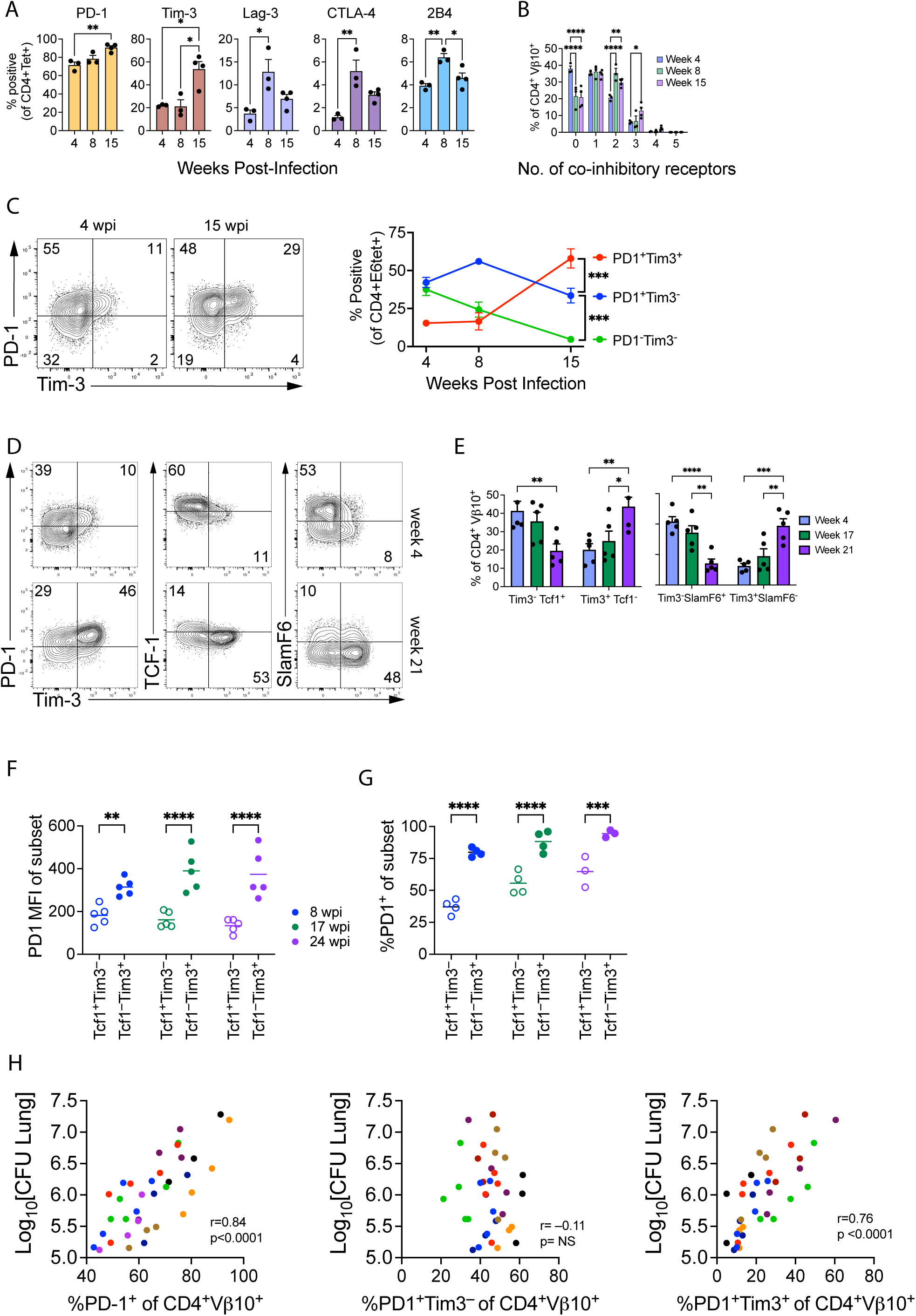
C7 T cells acquire an exhausted phenotype in the C7➔TCRα adoptive transfer model. (A) Frequencies of CD4^+^ ESAT-6 Tetramer^+^ C7 T cells expressing PD-1, TIM-3, Lag-3, CTLA-4, or 2B4. (B) Percentage of C7 T cells expressing 0 - 5 co-inhibitory receptors in the lungs of infected mice. (C) Representative flow plots and quantification of C7 T cells expressing PD-1 and/or TIM-3. (D) Representative flow plots of C7 T cells expressing TCF-1, SLAMF6, or PD-1, and TIM-3, at 4 and 21 wpi. (E) Frequencies of C7 T cells expressing TCF-1, SLAMF6, or TIM-3. (F, G) Percentage of TCF-1^+^TIM-3^-^ and TCF-1^-^TIM-3^+^ C7 T cells expressing PD-1 (F) and the PD-1 median fluorescence intensity (MFI) (G). (H) Correlation of lung CFU with percentage of PD-1^+^, PD-1^+^TIM-3^-^, or PD-1^+^TIM-3^+^ C7 T cells. Independent experiments, each with two or three timepoints are indicated by different colors. (A-G) Representative data of two (A, B) or more than three (C-G) independent experiments. (H) Cumulative data from 2 independent experiments. Data represent mean ± SEM. Statistical significance was analyzed by two-way ANOVA (E-G) with Bonferroni’s (A), Tukey’s (C, E), Fisher’s LSD multiple comparison test, or one-way ANOVA with Tukey’s multiple comparison test (B). Pearson correlation was calculated using data from three independent experiments consisting of 36 individual mice (H). p-values: *, p<0.05; **, p<0.01; ***, p<0.001; ****, p<0.0001; ns, no significant difference.

### Exhaustion contributes to CD4 T cell dysfunction in an adoptive transfer model

Based on the co-expression of PD-1 and TIM-3 by a large fraction of C7 CD4 T cells during the chronic phase of infection in this model, which resemble CD8 T_TEX_ cells, we hypothesized that C7 CD4 T cells were becoming exhausted ^11, 13^. To test this possibility, lung parenchymal CD4 T cells from C7➔TCRα mice were sorted (see Figure S1b), transcriptionally profiled at 4 and 25 wpi by RNAseq and compared to splenic CD4 T cells from uninfected C7 TCRtg mice (Figure 3a).

**Figure 3.**
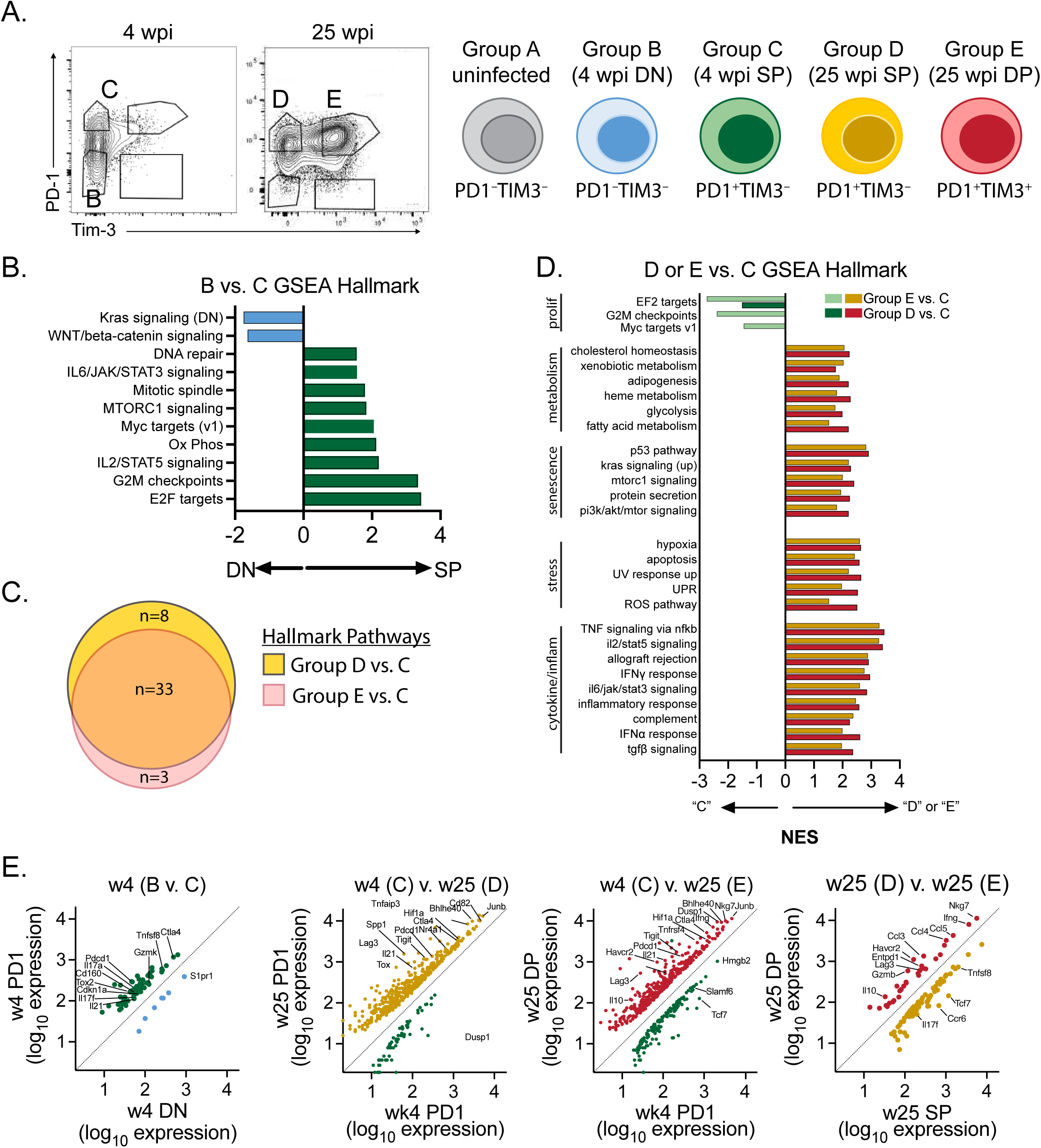
Exhaustion contributes to CD4 T cell dysfunction in the adoptive transfer model. (A) Flow plots and schematic of sorted cell populations at 4 and 25 wpi used for RNAseq analysis. (B) GSEA results of mouse-ortholog Hallmark pathways significantly enriched among Group B or Group C C7 T cells. (C) Venn diagram representing the overlap of Hallmark pathways significantly enriched in Groups D or E compared to Group C. (D) GSEA Hallmark pathways significantly enriched among Group D or Group E compared to Group C. (E) Scatter plots of DEGs (|log_2_FC| ≥ 1 and Padj < 0.05) between indicated groups. NES, Normalized Enrichment Score; False Discovery Rate (FDR), <0.05 for all pathways shown.

At 4 wpi, CD4 T cells from C7➔TCRα mice were mostly PD-1^-^TIM-3^-^ (Double Negative, DN; Group B) or PD-1^+^TIM-3^-^ (Single Positive; SP; Group C). Compared to CD4 T cells from uninfected C7 mice, 3,777 and 4,268 genes, respectively, were upregulated (log_2_FC≧1, FDR<0.05) (Table S1). Jun and Fos were among the most highly induced genes, and Gene Set Enrichment Analysis (GSEA) identified gene signatures associated with T cell activation in both Group B and C C7 CD4 T cells. Only 62 differentially expressed genes (DEGs) were identified between Group B and C cells (Table S2). Among these genes were Il17a, l17f, Il21 and Tox2, suggesting that the PD1^+^ C7 CD4 T cells contains Th17-and Tfh-like cells. The EF2 targets, G2M checkpoint, and IL-2/STAT5 signaling Hallmark Pathways were enriched among Group C cells, indicating that PD1^+^ C7 CD4 T cells undergo greater proliferation and activation than Group B cells (Figure 3b).

By 25 wpi, nearly all C7 CD4 T cells were PD1^hi^TIM3^-^ (Group D) or PD-1^hi^TIM-3^+^ (Double Positive; DP; Group E) (Figure 3a). Most Hallmark pathways that were significantly enriched were identified both in group D and E when compared to group C (Figure 3c, Table S3 and S4). The only Hallmark pathways upregulated in group C were “EF2 targets,” “G2M checkpoints” and “Myc targets v1,” consistent with the greater proliferative activity of CD4 T cells during the early phase of Mtb infection (Figure 3d). The sharing of upregulated pathways by Group D and E suggests that the difference in the duration of infection (i.e., 4 vs 25 wpi) was more important in determining the transcriptional program of the cells than their expression of co-inhibitory receptors (i.e., PD1^hi^TIM3^-^ vs. PD-1^hi^TIM-3^+^). Highly enriched signaling pathways in groups D and E included those induced by T cell cytokines (IL-2, IFNγ), inflammatory cytokines (IL-6, TNF, IFNα), and anti-inflammatory cytokines (TGFβ). Lipid and other metabolic pathways were also detected (fatty acid metabolism, cholesterol homeostasis, glycolysis). Genes from pathways associated with cellular stress were enriched in C7 CD4 T cells from 25 wpi including “apoptosis,” “hypoxia,” “ROS pathway,” “UPR,” and “UV response up,” which often indicate cell stress and DNA damage. The upregulation of these pathways could be driven by the infection and might contribute to T cell dysfunction.

Compared to Group C (4 wpi), 463 genes in Group D and 489 genes in Group E significantly changed expression, and 80% of these were up regulated at 25 wpi (Table S5 and S6). *Ccr4*, *Ccr6*, *Il17f*, *Il17re*, and *Il23r*, were expressed more at 4 wpi, suggesting that Th17 responses diminished during chronic infection. The expression of negative regulators and co-inhibitory receptors by Group D and E C7 CD4 T cells (PD1^hi^TIM3^-^ and PD1^hi^TIM3^+^, respectively) were increased (e.g., *Ctla4*, *Entpd1*, *Foxp3*, *Lag3*, *Nfil3*, *Pdcd1*, and *Tigit*) (Figure 3e, Table S5 and S6). *Tox* and *Tox2*, and *Nr4a1*, *Nr4a2*, and *Nr4a3*, are important transcriptional regulator of CD8 T cell exhaustion, and were increased among Group D and E C7 CD4 T cells. Among the Group E CD4 T cells, we noted increased expression of genes associated with CD4 T cell exhaustion (e.g., *Prdm1*, *Il10*, *Il21*) ^23^ and terminal differentiation (*Klrg1* and *Tbx21*). Thus, both 25 wpi CD4 T cell populations had features resembling exhausted CD8 T cells observed in tumors and chronic viral infection ^13, 24, 25^.

Only 108 genes differed between 25wpi PD1^hi^TIM3^-^ and 25wpi PD1^hi^TIM3^+^ CD4 T cells (Figure 3e, Table S7). Like the 4 vs. 25 wpi comparison, Group E CD4 T cells not only expressed higher levels of genes associated with exhaustion (*Havcr2*, *Entpd1*, *Lag3*, *Il10*) but also soluble mediators (*Ifng*, *Ccl3*, *Ccl4*, *Ccl5*) and genes associated with a cytolytic phenotype (*Gzmb*, *Nkg7*, *Prf1*). In contrast, Group D CD4 T cells expressed higher levels of *Slamf6* and *Tcf7*, indicating a similarity to CD8 T_PEX_ cells. We conclude that in the C7➔TCRα model, C7 CD4 T cells acquire a transcriptional signature during the chronic phase of infection that resembles exhausted CD8 T cells.

### Senescence signature among CD4 T cells during chronic infection

Three Hallmark pathways that were significantly enriched during chronic infection (i.e., expressed in Group D and E, compared to Group C C7 CD4 T cells) were the “p53 pathway,” “Hypoxia,” and “Interferon alpha response” gene sets (Figure 3d, 4a). p53 is the best studied human tumor suppressor gene and is the most frequently mutated gene among human cancers. However, physiologically, the p53 pathway orchestrates key responses, often in response to DNA damage, oxidative damage, or metabolic stress, and generates a variety of cellular responses including DNA repair, cell cycle arrest, apoptosis, and senescence ^26^. Activation of the p53 pathway has recently been linked to T cell senescence ^27, 28^. Both Type I IFN and hypoxia can also lead to cellular senescence (Figure 4a) ^29–31^.

**Figure 4.**
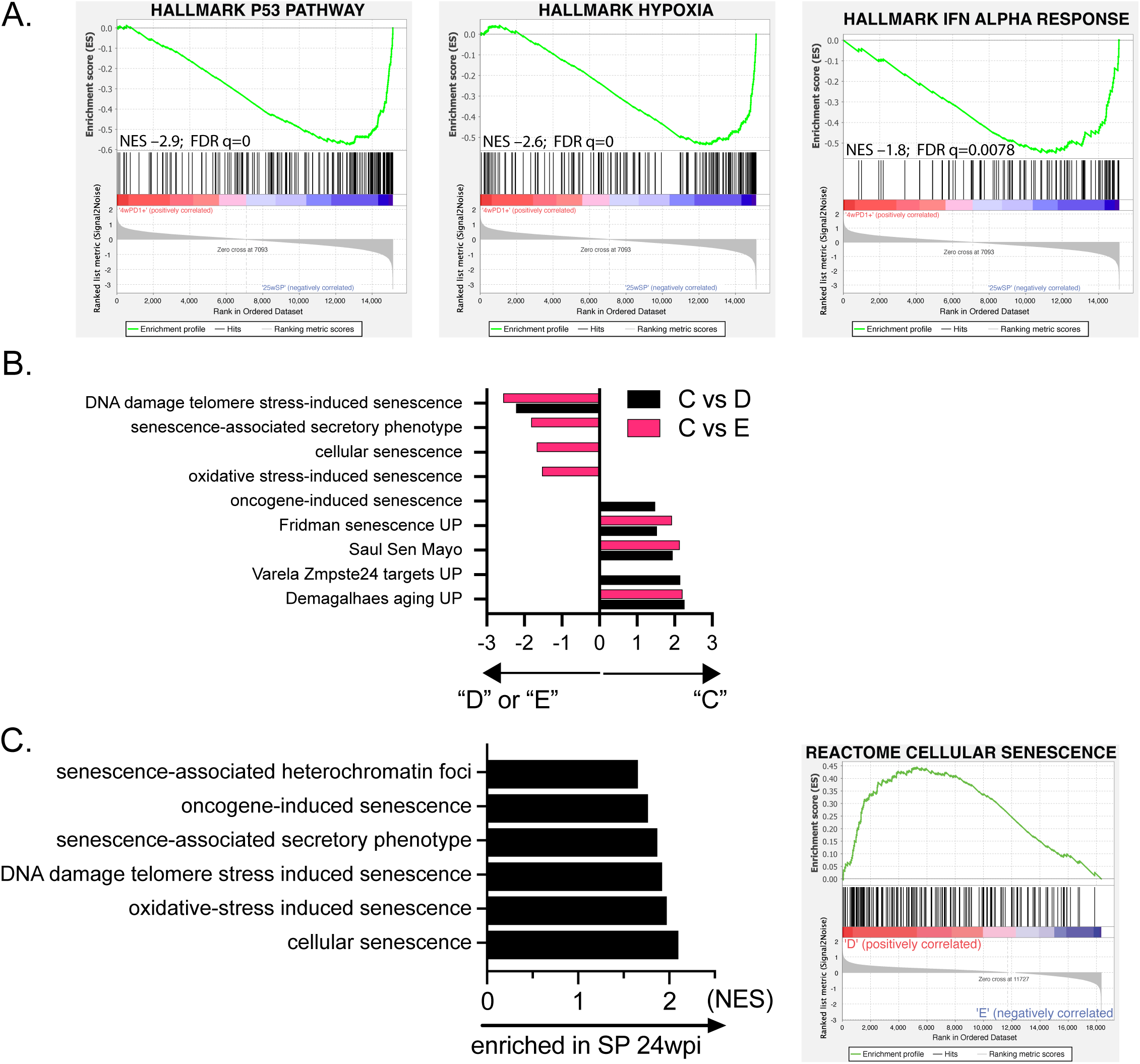
Senescence-related signatures were detected among C7 CD4 T cells during chronic infection. (A) GSEA Enrichment plots between Group C and Group D T cells for the p53, Hypoxia, and IFN alpha Hallmark pathways. (B) GSEA results of senescence-related signatures enriched in Groups D or E compared to Group C. X axis, NES. (C) Senescence pathways enriched in Group D compared to Group E (left) and the Cellular Senescence GSEA enrichment plot (right). FDR, <0.05 for all pathways shown.

We hypothesized that transcriptional pathways in response to cell stressors (UV damage, hypoxia, apoptosis, UPR, ROS pathway) could be mechanistically linked by activating the p53 pathway. Therefore, we considered whether the Hallmark pathways of “protein secretion,” “Mtorc1” and “PI3K/AKT3/mTOR” signaling, cytokine signaling (IFNγ, IFNα, TNF, TGFβ, IL-6) during chronic infection could indicate CD4 T cell senescence (Figure 3d). Analysis using a compendium of “Senescence” signatures (Table S8) revealed significant enrichment of the ‘Saul Sen Mayo’ and ‘Fridman Senescence Up’, and other gene sets ^32^, which indicate that a program of cell senescence is active late during infection (Figure 4b). Group D and E cells differed with respect to transcriptional programs associated with senescence. Several senescent gene sets were significantly enriched in Group D (PD1^hi^TIM3^-^) but not Group E (PD1^hi^TIM3^+^) C7 CD4 T cells (Figure 4c). Also, PD-1 and TIGIT are associated with senescence and aging ^33, 34^. p53 and p16 were the two genes most frequently shared in a leading-edge analysis. Thus, while chronic infection leads to changes consistent with senescence and exhaustion in C7 CD4 T cells, PD1^hi^TIM3^-^ C7 cells (Group D) are more typical of senescent T cells while PD1^hi^TIM3^+^ C7 cells (Group D) represent exhausted T cells.

### Few lung-resident CD4 T cells express multiple effector function during chronic infection

Given the surprising heterogeneity of CD4 T cell responses, especially regarding the detection of both effector and dysfunctional features, we next assessed whether the apparent heterogeneity could be resolved using scRNAseq to analyze polyclonal CD4 T cell responses. We investigated whether CD4 T cells become dysfunctional during chronic Mtb infection by comparing CD4 T cells from the lungs of C57BL/6 (B6) mice early (4 - 6 wpi) vs. late (39 - 50 wpi) after Mtb infection. CD4^+^CD44^-^ T cells expressed little PD-1 or TIM-3, consistent with these cells being naïve (Figure 5a). Early after infection, most CD44^+^E6^-^ CD4 T cells (antigen-experienced excluding ESAT-6-specific) expressed PD-1 and a smaller population also expressed TIM-3. More ESAT-6 specific CD4 T cells (i.e., CD44^+^E6^+^tet) expressed both PD-1 and TIM-3. In contrast, nearly all CD4 T cells expressed PD-1 during chronic infection. Importantly, the ESAT6-specific CD4 T cells were predominantly PD-1^+^TIM-3^+^ late during chronic infection (Figure 5a). Thus, CD4 T cells in chronically infected mice express both PD-1 and TIM-3, a phenotype associated with CD8 T_TEX_ cells ^11, 13^.

**Figure 5.**
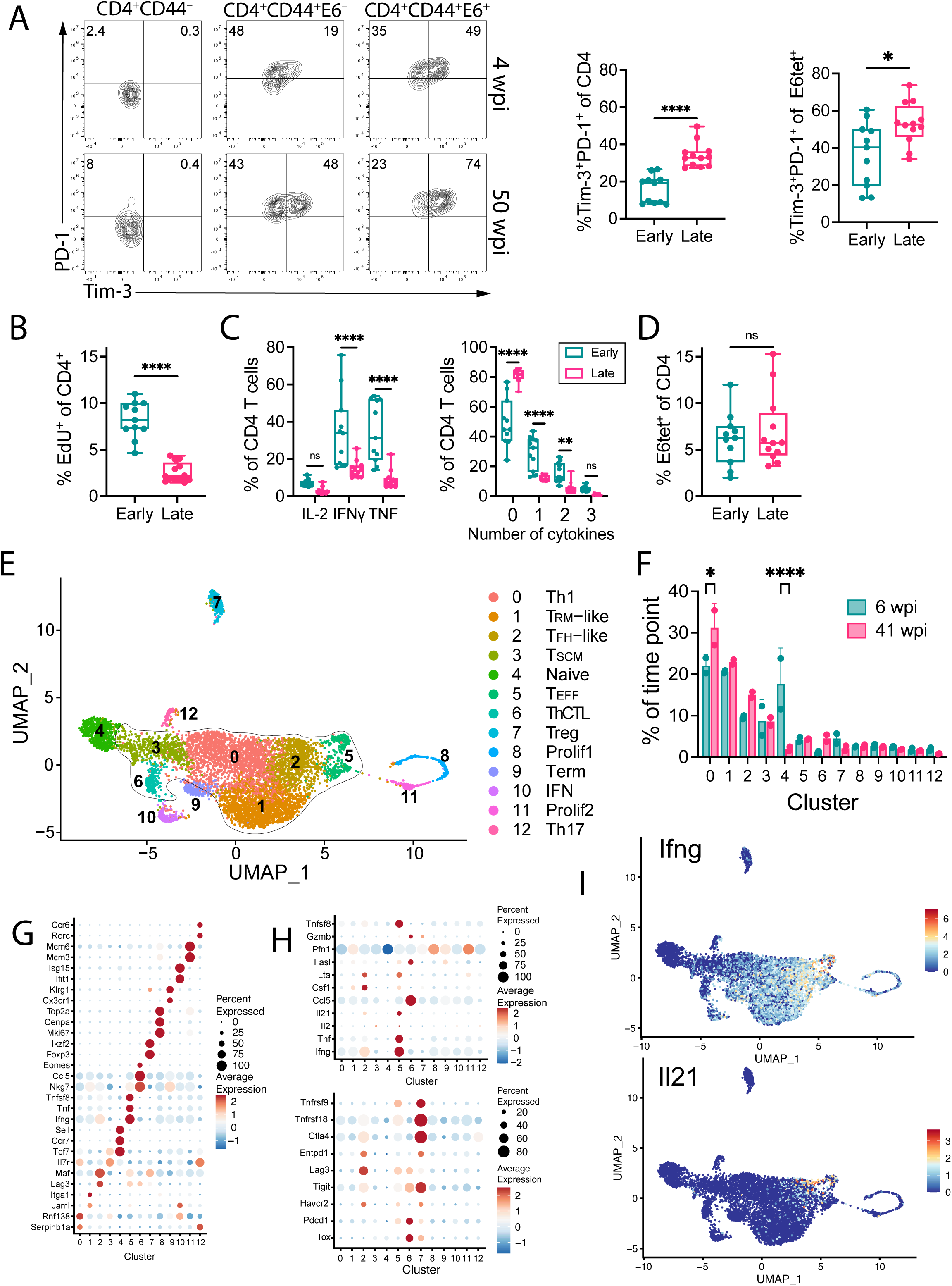
Few lung-resident CD4 T cells express multiple effector function during chronic infection. (A-D) Data is from 24 mice in 3 experiments infected for 4 – 6 (early), or 39 – 50 (late) weeks. E6tet^+^, Esat-6 tetramer^+^. (A) CD4 T cells express both PD-1 and TIM-3 during chronic infection. (B) Percentage of EdU^+^ CD4^+^CD44^+^ T cells early (4 - 6 wpi) or late (39 - 50 wpi) after infection. (C) Percentage of CD4^+^CD44^+^ T cells producing IL-2, IFNγ, or TNF, after stimulation with ESAT6_1-15_ (E6-stim). (D) Percentage of ESAT6-tetramer^+^ (E6tet^+^) CD4^+^CD44^+^ T cells among total CD4 T cells, early or late after infection. (E-I) Total lung parenchymal T cells were sorted and analyzed by scRNAseq. (E) UMAP projection of CD4 T cells clustered based on scRNAseq analysis and cluster designations. Outlined clusters all have a Th1 core signature. (F) Distribution of CD4 T cells among the different clusters, normalized by the total number of CD4 T cells from each subject. (G) Dot plot of differentially expressed genes. (H) Dot plot of genes associated with effector functions or co-inhibitory receptors. (I) UMAP projections overlapped with expression of *Ifng* and *Il21*.

We next assessed the function of CD4 T cells during chronic infection. Significantly fewer cells proliferated compared to early timepoints (Figure 5b). Late during infection, fewer CD4 T cells produced IFNγ and TNF after ESAT6 stimulation, and polyfunctionality was lost over time (Figure 5c). Importantly, the frequency of ESAT-6 specific CD4 T cells was similar at both timepoints post-infection; thus, differences were not due to altered numbers of ESAT6-specific CD4 T cells (Figure 5d). These data show that activated CD4 T cells develop a dysfunctional T cell phenotype during chronic infection.

To characterize CD4 T cell heterogeneity during chronic infection, CD4 IV^-ve^ parenchymal lung T cells were flow sorted from Mtb-infected mice 6 and 41 wpi for analysis by scRNAseq and TCRseq (Figure S2a). Cell clustering based on gene expression (excluding TCR genes) identified 13 clusters (Figure 5e). Only the proportion of Cluster 0 and Cluster 4 differed between 6 and 41 wpi (Figure 5f, Figure S2b). The CD4 clusters were annotated using transcriptional signatures and key canonical genes (Figure 5g-h; Figure S2c, Table 1, Table S9). Clusters 7 – 12 accounted for 13% of the CD4 cells. Cluster 7 were Tregs (*FoxP3, Ctla4, IL2Ra, Ikzf2*). Cluster 8 and 11 were proliferating cells (*Mki67* and *Top2a* in G2/M or G1/S cell cycle phase, respectively). Cluster 9 expressed high levels of *Klrd1*, *Klrg1*, *Tbx21* (T-bet) and *Cx3cr1*, indicating that these were terminally differentiated CD4 T cells. Cluster 10 expressed a type I IFN signature (*Isg15, Bst2, Ifit1, Ifit2 and Ifit3*). Cluster 12 were Th17 cells (*Ccr4, Ccr6, 1l17re, Rorc, Aqp3*).

**Table 1.** CD4 T cell clusters.

| **Cluster** | **Description** | **Upregulated genes** | **Core Th1** |
| --- | --- | --- | --- |
| 0 | Th1 | *Cxcr3, Gzmk, Icos, Ifngr1, Il18r1* | yes |
| 1 | TRM-like | *Cd44, Itga1, Itgb2, Jaml* | yes |
| 2 | TFH-like | *Icos, Ifng, Il21, Lag3* | yes |
| 3 | TSCM | *Il7r, S1pr1, Tcf7* | yes |
| 4 | Naive | *Ccr7, Lef1, Sell, Tcf7* | – |
| 5 | TEFF | *Ccl3, Ifng, Il21, Lta, Tnf, Tnfsf8* | yes |
| 6 | ThCTL | *Eomes, Nkg7, Tox* | yes |
| 7 | TREG | *Ctla4, Foxp3, Havcr2, Il10, Tgfb1* | – |
| 8 | Prolif1 (G1/S) | *Mki67, Top2a, Tuba1b* | – |
| 9 | Terminal diff. | *Ccl5, Cx3cr1, Klrg1, Nkg7, Tbx21* | – |
| 10 | Type I IFN | *Bst2, Ifit1, Ifit2, Ifit3, Irf7, Isg20* | – |
| 11 | Prolif2 (G2/M) | *Mki67, Top2a, Tuba1b* | – |
| 12 | Th17 | *Ahr, Ccr4, Ccr6, Il17a, Il17f, Rorc* | – |

Six clusters, including the four largest clusters (0, 1, 2, 3), all expressed *Cxcr3, Cxcr6, Id2, Nkg7, Ccl5 and Ifng,* suggesting that they were different forms of Th1 cells (Figure 5e). We used FindMarkers to compare Clusters 0, 1, 2, 3, 5 and 6, to naïve-like CD4 T cells (Cluster 4; *Crr7*, *Sell, Lef1*) (Table S9). Cluster 6 cells expressed high levels of *Ccl3, Ccl4*, *Gzmb* and *Eomes*; some cells expressed *Pfn1*. As Eomes promotes a cytotoxic program in human CD4 T cells, Cluster 6 could be cytotoxic CD4 T cells ^35, 36^. Clusters 0, 1, and 3 were difficult to annotate. Cluster 0 resembled Th1 cells (*Cxcr3, Cxcr6, Id2, Nkg7, Ccl5, Ifng)* and Cluster 1 expressed genes indicative of tissue residency (*Itga1, Jaml, CD226*)^37, 38^. Finally, upregulated genes in Cluster 3 included *Tcf7, Il7r, Il2rb, Ly6a, Cxcr3, S1pr1*, and *Klf2* suggesting these are TSCM ^39^. *Ifng* transcript was detected in Clusters 0, 1, 2, 3, 5 and 6, but Clusters 2 and 5 expressed considerably more *Ifng* than the other clusters. Cluster 5 cells expressed multiple effector genes required for protection including *Tnf, Il21, Lta,* and *Tnfsf8* (Figure 5h; Table S9) ^19, 40, 41^. CD4 T cells belonging to Cluster 5 are the key polyfunctional effector CD4 T cells in the lungs of infected mice. Cluster 2, which is adjacent to Cluster 5, also expressed *Ifng, tnf, Il21,* and *Tnfsf8,* but lower amounts (Figure 5h). In chronically infected B6 mice, many parenchymal CD4 T cells had a PD-1^+^TIM-3^+^TCF7^-^ phenotype based on flow cytometry, which is typical of CD8 T_TEX_ cells ^11, 13^. However, this cell surface phenotype poorly correlated with the transcriptional signature apart from Cluster 2, which we believe are exhausted. Notably, when the DEGs from the C7 RNAseq were used as a gene signature to probe the B6 scRNAseq dataset, the genes upregulated in Group D and E (vs. Group C, Figure 3e) were most highly expressed by Cluster 2 and 5. The DEGs upregulated by Group E (vs. Group D) were specifically expressed by Cluster 2, indicating that these polyclonal CD4 T cells resemble exhausted T cells (Figure S2d).

Finally, polyclonal CD4 T cells in chronically infected B6 mice also expressed p53, senescence, and hypoxia signatures, particularly within Clusters 0, 1, 2, and 5 (Figure S3a-d). Thus, it appears that Cluster 5, which is only 3-4% of the CD4 T cells, are the true polyfunctional effectors that express the highest levels of IFNγ, IL-21, and TNF (Figure 5I). In contrast, the majority of other Th1 cells appear to be hypofunctional, especially with respect to cytokine production.

### Early and late CD4 T cell responses express distinct gene signatures

We identified statistically significant DEGs that were upregulated at 6 or 41 wpi using Seurat FindMarkers (Table S10). Panther was used to determine molecular pathways that were statistically over-represented in the input list compared with randomly selected genes^42^. Annotated pathways including “T cell activation (P00053),” “JAK/STAT signaling (P00038)”, “Hypoxia response via HIF activation (P00030),” and “Inflammation mediated by chemokine and cytokine signaling (P00031),” were predicted to be upregulated at 6 wpi. No pathways of immunological relevance were significantly upregulated at 41 wpi. Next, genes that were upregulated at 6 or 41 wpi in each cluster were used to query the Reactome database (Table S11). We focused on Clusters 0, 1, 2, 3, and 5, since they were of greatest interest. These five clusters were enriched for TCR signaling and CD28 costimulation pathways (except Cluster 3) at 6 wpi (Table 2). Clusters 0, 2, and 5 appeared most activated based the identification of multiple signaling pathways including those for interferon, IL-2, and IL-21 signaling (Table 2, S11). At 41 wpi, only Clusters 2 and 3 were enriched for TCR signaling and none of the clusters were enriched for cytokine signaling pathways.

**Table 2.** Panther analysis of pathways enriched early (6 wpi) or late (41 wpi) during infection in each cluster.

|  | Th1 |  | TRM-like |  | TFH |  | TSCM |  | Teff |  |
| --- | --- | --- | --- | --- | --- | --- | --- | --- | --- | --- |
|  | Cluster 0 |  | Cluster 1 |  | Cluster 2 |  | Cluster 3 |  | Cluster 5 |  |
|  | 6 wpi | 41 wpi | 6 wpi | 41 wpi | 6 wpi | 41 wpi | 6 wpi | 41 wpi | 6 wpi | 41 wpi |
| REACTOME PATHWAYS |  |  |  |  |  |  |  |  |  |  |
| Regulation of gene expression by HIF (R-MMU-1234158) | 6.53E-03* |  | 4.88E-03 |  | 7.85E-03 |  | 7.72E-03 |  | 6.68E-03 |  |
| Interferon Signaling (R-MMU-913531) | 8.21E-03 |  | 4.68E-03 |  | 6.38E-03 |  | 4.82E-03 |  | 8.85E-03 |  |
| TCR signaling (R-MMU-202403) | 8.70E-06 |  | 1.34E-05 |  | 1.58E-04 | 2.09E-03 | 2.38E-04 | 4.06E-03 | 7.37E-06 |  |
| Downstream TCR signaling (R-MMU-202424) | 8.70E-06 |  | 1.84E-02 |  | 4.89E-02 | 8.62E-04 |  | 8.62E-04 | 9.40E-03 |  |
| CD28 co-stimulation (R-MMU-389356) | 1.61E-03 |  | 2.88E-02 |  | 1.15E-02 |  |  |  | 4.28E-02 |  |
| TNF signaling (R-MMU-75893) | 1.27E-02 |  |  |  | 5.41E-03 |  |  |  | 4.73E-02 |  |
| IL-2 family signaling (R-MMU-451927) | 4.13E-03 |  |  |  | 6.07E-06 |  |  |  | 3.77E-03 |  |
| IL-2 signaling (R-MMU-9020558) |  |  |  |  | 3.34E-03 |  |  |  |  |  |
| IL-21 signaling (R-MMU-9020958) |  |  |  |  | 1.95E-03 |  |  |  |  |  |
| Regulation of NF-kappa B signaling (R-MMU-9758274) |  |  |  |  | 4.37E-02 |  |  |  | 3.47E-02 |  |
| Cellular Senescence (R-MMU-2559583) | 4.07E-02 |  |  |  | 1.19E-02 |  | 4.07E-02 |  | 1.77E-02 |  |
| Oxidative Stress Induced Senescence (R-MMU-2559580) | 1.31E-02 |  | 1.58E-03 |  | 8.57E-04 |  | 1.73E-02 |  | 2.49E-03 |  |
| Signaling by TGFB Receptor Complex (R-MMU-170834) | 2.94E-04 |  | 3.77E-06 |  | 1.19E-06 |  | 4.88E-07 |  | 2.14E-03 |  |
| Signaling by TGFB family members (R-MMU-9006936) | 1.86E-03 |  | 5.86E-05 |  | 2.74E-05 |  | 9.85E-06 |  | 2.96E-05 |  |
| TGFB receptor signaling activates SMADs (R-MMU-2173789) |  |  | 4.20E-05 |  | 2.90E-05 |  | 1.34E-05 |  | 8.36E-05 |  |
| Downregulation of TGFB receptor signaling (R-MMU-2173788) | 5.57E-03 |  | 4.03E-03 |  | 7.56E-03 |  | 7.27E-03 |  | 3.18E-02 |  |
| CTLA4 inhibitory signaling (R-MMU-389513) | 1.24E-02 |  | 1.19E-03 |  | 1.70E-02 |  | 2.55E-04 |  |  |  |
| IL-27 signaling (R-MMU-9020956) |  |  | 1.41E-02 |  | 1.41E-02 |  | 2.07E-02 |  | 1.91E-02 |  |
| Transcriptional Regulation by TP53 (R-MMU-3700989) | 2.35E-04 |  | 1.99E-04 |  | 2.08E-03 |  | 6.11E-06 |  | 1.29E-03 |  |
| Signaling by VEGF (R-MMU-194138) | 1.69E-05 |  | 6.89E-04 |  | 6.48E-05 |  | 4.19E-04 |  | 4.08E-03 |  |
| Transcriptional regulation by RUNX1 (R-MMU-8878171) | 3.65E-04 |  | 1.58E-04 |  | 9.64E-04 | 2.55E-03 | 1.60E-04 |  | 2.67E-05 |  |
| Signaling by WNT (R-MMU-195721) | 1.33E-05 |  | 1.58E-04 |  | 7.65E-03 | 7.65E-03 | 2.72E-05 |  | 1.00E-03 |  |
| Signaling by NOTCH1 (R-MMU-1980143) | 1.53E-03 |  | 1.03E-03 |  | 2.19E-03 |  |  |  | 1.46E-03 |  |

\* Values indicate FDR. Blanks indicate FDR>0.05. See Table S11 for all significant pathways.

Regulatory pathways including CTLA4 signaling and signaling by the IL-27 and TGFβ inhibitory cytokines were enriched at 6 wpi but not 41 wpi. TGFβ signaling was confirmed by applying a TGFβ signaling signature to the UMAP projection (Figure 6a). Genes induced by TGFβ were upregulated in several clusters at 6 wpi, but by 41 wpi, the signature was prominently expressed only by CD4 T cells in Cluster 1. Consistent with these data, *Tgfbr2* was more highly expressed at 6 wpi than 41 wpi (Figure 6b). Interestingly, genes in the “Downregulation of TGFB receptor signaling (R-MMU-2173788),” pathway were also significantly overrepresented at 6 wpi, suggesting that TGFβ receptor signaling was being actively regulated early but not late during the immune response to Mtb. Pathway pertaining to Notch and Runx signaling, which can synergize with other transcription factors to promote T cell activation and Th1 differentation, were enriched in several clusters at 6 wpi but not 41 wpi (Table S11). Genes upregulated in several clusters at 6 wpi, and to a more limited degree at 41 wpi, were enriched in pathways pertaining to cell stress and injury including senescence, apoptosis and necrosis, and hypoxia (Table 1, Table S11). Differences in individual genes were detected between 6 and 41 wpi. Various combinations of *Ifng, Ccl5, Tnfsf8, Gzmk, Nkg7,* and *Il7r* were significantly increased in Clusters 0, 1, 2, and 5 at 41 wpi (Figure 6c, Table S10). We speculate that the increased expression of these genes at 41 wpi could reflect an alteration of the regulatory environment of the Mtb lesion, or a consequence of greater antigen load.

**Figure 6.**
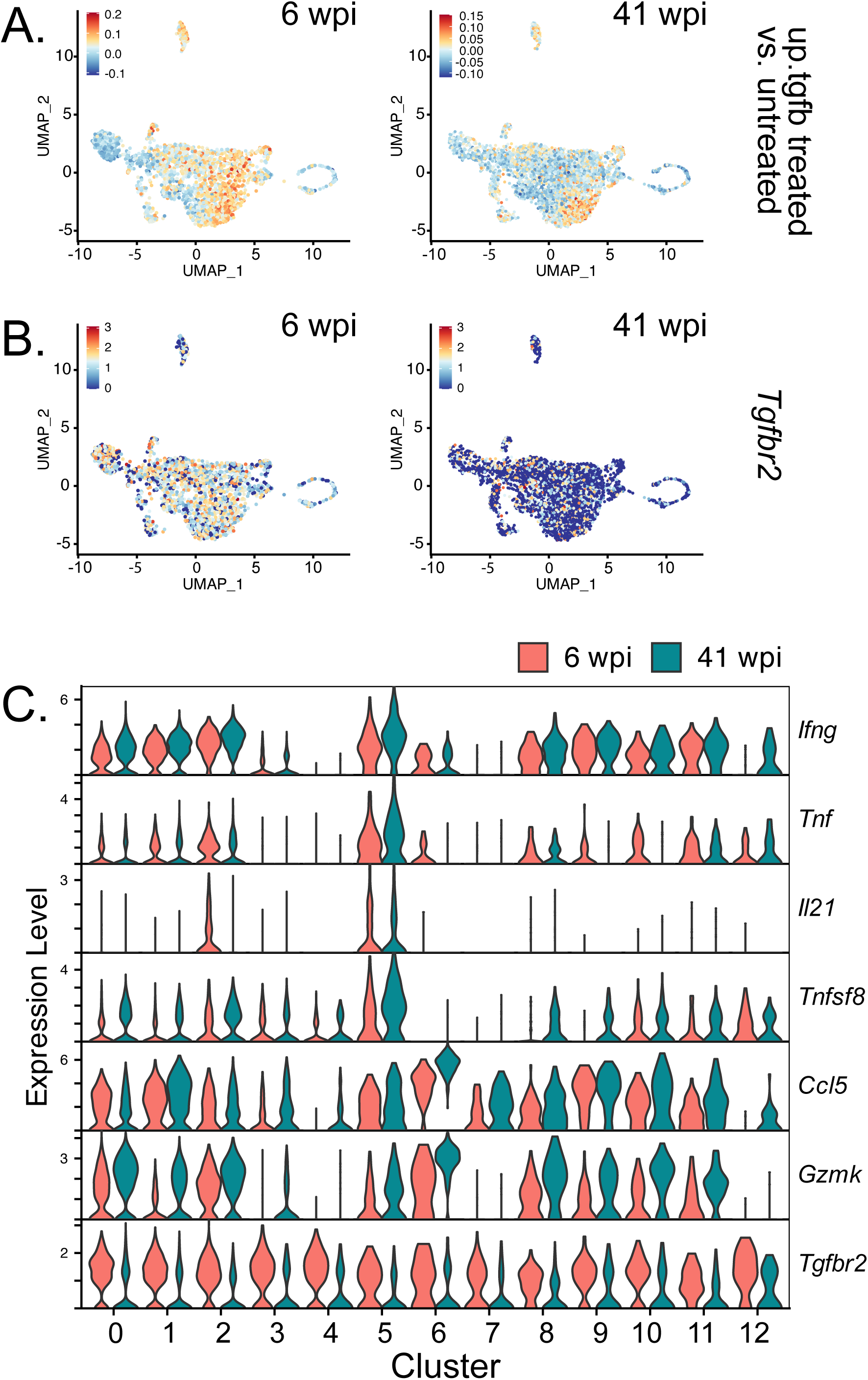
Transcriptional differences between CD4 T cells in the lungs of mice, early (6 wpi) or late (41 wpi) after infection. (A). UMAP projections overlapped with expression of *Tgfb* signaling signature. (B) UMAP projections overlapped with expression of *Tgfrb2* gene expression. (C) Violin plots of selected genes that show differential expression between 6 wpi vs. 41 wpi.

Increased expression of these effectors could drive more inflammation or even be a sign of CD4 T cell dysfunction.

### Large clonal expansions of activated CD4 T cells are maintained during chronic infection

TCR analysis was used to identify CD4 T cells responding to Mtb since clonal expansion following activation is a cardinal feature of antigen-specific T cell responses. We identified 5,573 paired TCRαβ; 2,321 from 6 wpi and 3,252 from 41 wpi. There were 1,965 unique clonotypes based on amino acid sequences, which were categorized by the size of the expansion (Figure 7a, 7b). The majority of TCRs in all clusters belonged to highly expanded clonotypes (i.e., >5), except for Clusters 3, 4, 7, and 12. In Cluster 4, 99% of the TCR clonotypes were singletons, consistent with their annotation as naïve CD4 T cells. The 10 most abundant clonotypes accounted for 21% of CD4 T cells. We assessed the top 10 clonotypes for each subject (Figure 7c). More than 75% of these highly expanded clonotypes had a TH1, TRM-like, TFH-like, or TEFF phenotype (i.e. clusters 0, 1, 2, 5, respectively) and were largely absent from the naïve, TREG and TH17 clusters. Overall, the TH1, TRM-like, and TFH-like clusters were dominated by highly expanded clonotypes (i.e., containing >20 cells), which accounted for 47-63% of the TCRs in these clusters. As a control for antigen-specificity, C7 CD4 T cells from uninfected mice were transferred into each B6 mouse one day before their infection. The C7 cells were identified based on their CDR3β sequence (Table S12) ^43^. Like endogenous CD4 T cells, most C7 cells were in the TH1, TRM-like, TFH-like, or TEFF clusters (Table S12). These data show that while public TCRs for Mtb antigens exist and convergent recombination occurs, most clonal expansions were derived from a single CD4 T cell. Importantly, the detection of Mtb-specific CD4 T cells from the same clonotype among different transcriptional clusters implies that their differentiation and activation is influenced by extrinsic factors in microenvironments of the infected lung. These data show that Mtb infection is driving CD4 T cell responses in these clusters.

**Figure 7.**
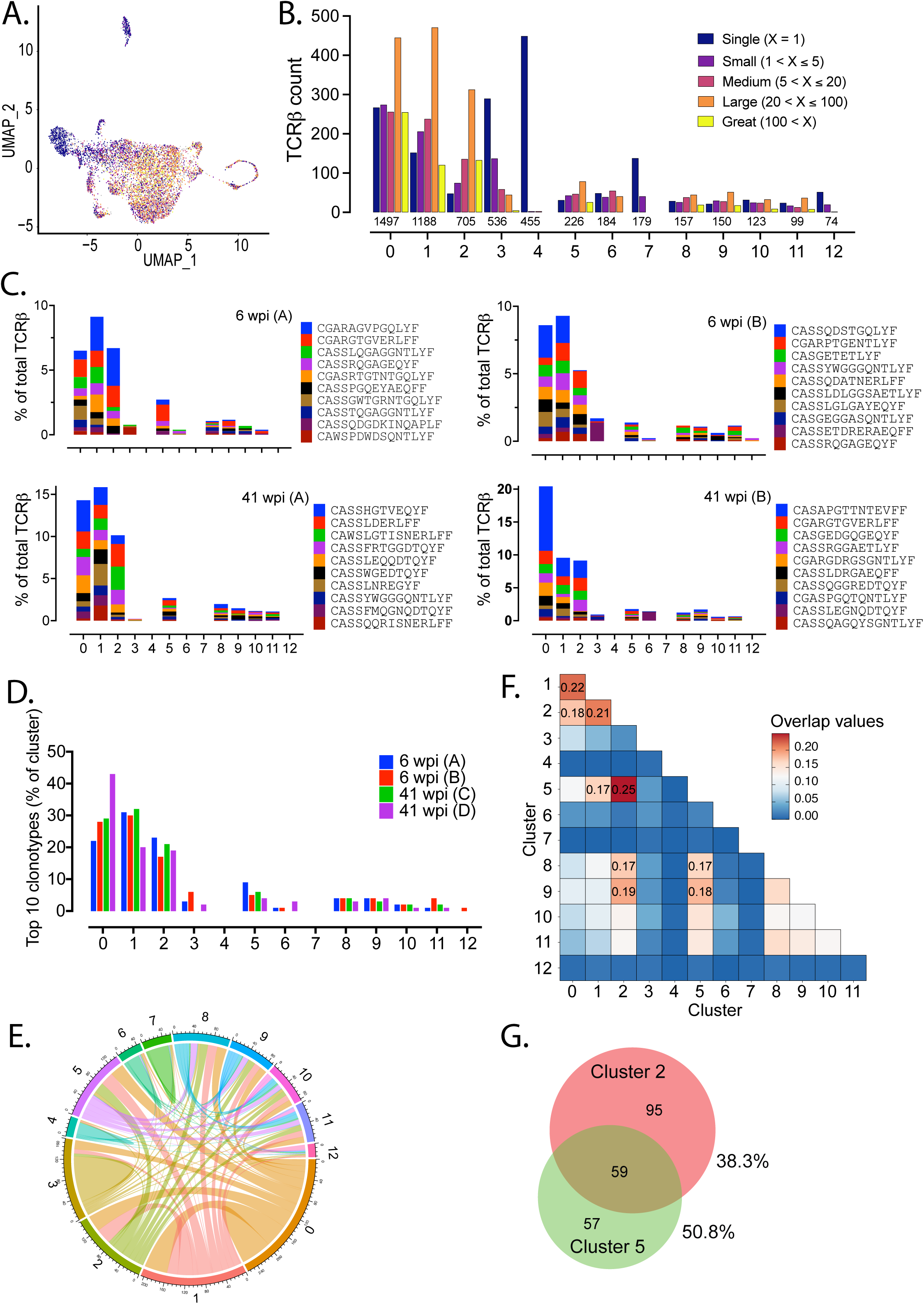
Large clonal expansions of activated CD4 T cells accumulate in Clusters 0, 1, and 2, and are maintained during chronic infection. (A) A UMAP projection colored by the size of the TCR clonotype expansions. (B) Distribution of TCR count and clonotype size in each CD4 cluster. The number of total TCRs is listed above each cluster designation. (C) For each subject analyzed by TCRseq, the distribution of the 10 most abundant CDR3β among the 13 clusters is shown. (D) For each subject, the percentage of the total TCRs in each cluster accounted for by the top 10 TCR clonotypes. (E) A chord diagram of shared TCRs between clusters for Mouse 1 (46 wpi, A). (F) The overlap of TCR clonotypes between clusters based on the Jaccard Similarity Index. (G) Sharing of clonotypes between Clusters 2 and 5 determined by Venn analysis ^89^.

Many of the highly expanded clonotypes were distributed among several clusters (Figure 2e). To quantify the degree of TCR sharing, we calculated the Jaccard Similarity Index between each cluster pair. Significant sharing was detected between Clusters 0, 1, 2, and 5 (Figure 7f). The greatest amount of TCR sharing was between Cluster 2 and 5, which were adjacent to each other on the UMAP projection and shared some transcriptional characteristics (Figure 5). Sharing of unique clonotypes for Cluster 2 and Cluster 5 was 38.3% and 50.8%, respectively (Figure 7g). Cluster 5 had the greatest expression of genes associated with CD4 effector function. Finding that Cluster 0, 1, and 2 CD4 T cells, highly related by TCR clonotype and parenchymal location to Cluster 5 TEFF cells, did not express any effector T cell transcriptional program, suggests these cells were hypofunctional or possibly paralyzed ^44^.

### TCR affinity affects the development of exhaustion

As TCR affinity and signaling strength affect T cell differentiation ^45–48^, we hypothesized that TCR affinity would also influence the development of T cell dysfunction. C24 TCRtg mice express a TCR that also recognizes ESAT6_4-17_ as do C7 cells, but with higher affinity ^49^. C7 and C24 were transferred at a 1:1 ratio into TCRα KO mice and assessed at 4 and 34 wpi (Figure S1c). Early post-infection, no consistent phenotypic differences were observed between the two T cell clonotypes. At 34 wpi, the number of C7 and C24 cells was similar (Figure S1d). However, more C7 T cells resembled progenitor exhausted CD8^+^ T cells (i.e., TCF-1^+^TIM-3^-^), and more C24 T cells resembled terminally exhausted CD8^+^ T cells (i.e., PD-1^+^TIM-3^+^) (Figure 8a). C24 cells expressed more T-bet (Figure 8b), and more produced IFNγ (Figure 8c), indicating a more terminally differentiated phenotype. There were no differences in IL-2 or TNF production (Figure 8c).

**Figure 8.**
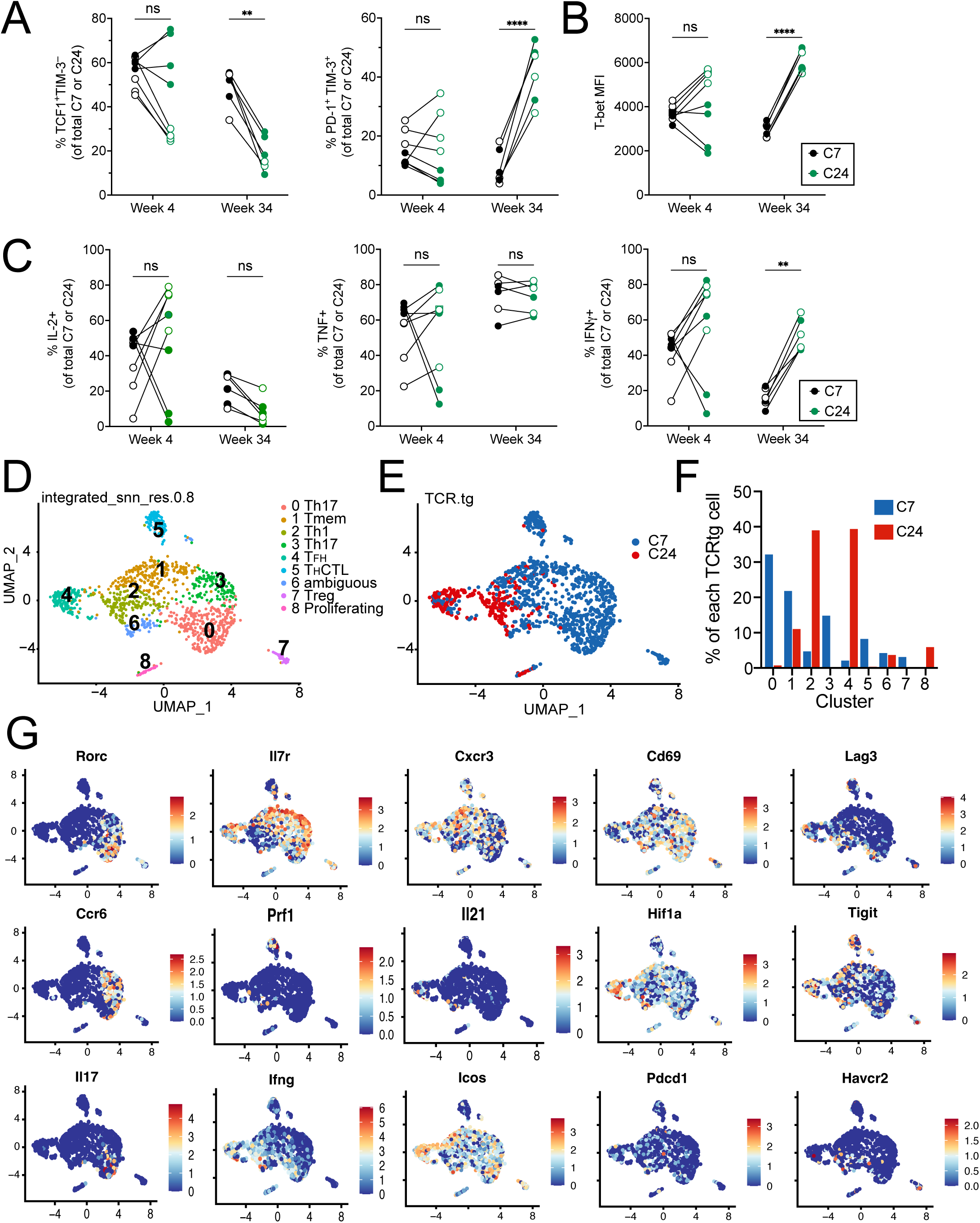
TCR affinity affects T cell function and differentiation in the adoptive transfer model of chronic Mtb infection. C7 (black) and C24 (green) TCRtg cells were co-transferred (1:1 ratio) into TCRα KO mice. Closed circles, females; open circles, males. Parenchymal C7 and C24 cells were analyzed at 4 and 34 wpi. (A) Percentage of C7 or C24 cells that expressed TCF-1^+^TIM-3^-^ (left) or PD-1^+^TIM-3^+^ (right). (B) T-bet MFI. (C) Percentage of C7 or C24 cells that produced of IL-2, TNF, or IFNγ after ESAT6 peptide stimulation in vitro. Data are representative of two independent experiments. Statistical significance was analyzed by two-way ANOVA. p-values: *, p<0.05; **, p<0.01; ***, p<0.001; ****, p<0.0001; ns, no significant difference. (D-G) scRNAseq was performed on parenchymal CD4 T cells after co-transferred C7 and C24 cells, 24 wpi. (D) UMAP projection of C7 and C24 cells clustered based on scRNAseq analysis. (E) Location of C7 and C24 cells in the UMAP projection based on TCR sequence. (F) Proportions of C7 or C24 T cells within each cluster. (G) Feature plots of genes of referred to in the text.

We next performed scRNAseq after co-transferred of C7 and C24 cells, 24 wpi. We identified 1,181 C7 and C24 cells based on their CDR3β sequences, which were clustered into 9 populations (Figure 8d, Table S13). Clusters 2, 4, and 8 were predominantly C24 T cells; Clusters 0, 1, 3, 5, and 7 were predominantly C7 T cells; and Cluster 6 was mixed (Figure 8e, 8f). Clusters 0 and 3 were Th17 cells (*Rorc, Ccr6*), although only Cluster 0 expressed *Il17a* (Figure 8g). Cluster 1 had a memory phenotype (*Il7r*, *Tcf7*, *Klf2*); Cluster 5 had a cytotoxic signature (*Gzmb, Prf1*); Cluster 7 were Tregs (*Foxp3*); and Cluster 8 were proliferating cells (*Mki67*). Clusters 2, 4, and 6 contained *Tbx21* expressing cells, but only Clusters 2 and 4 significantly expressed *Ifng*. Cluster 4 also expressed *Il21*. These results align with more IFNγ production by C24 cells. The reduced effector capacity of Cluster 2 in the context of low *Il7r* and high *Hif1a* suggests chronic stimulation^50^ or hypoxia ^51, 52^. These two clusters are reminiscent of Clusters 2 and 5 in B6 mice. In that analysis, Cluster 5 were highly functional effector CD4 T cells, and Cluster 2 consisted of CD4 T cells that had lost expression of some, but not all, effector genes (Figure 5). The identity of Cluster 6, which mostly consisted of C7 T cells, was ambiguous. These cells expressed high levels of genes associated with CD8 T cell exhaustion (*Nr4a1, Nr4a3, Egr2*), suggesting Cluster 6 contained exhausted T cells ^53, 54^.

Our findings align with studies showing that high affinity TCR clones skew towards a Th1 phenotype ^45^, establishing that TCR signaling strength contributes to the varied differentiation states of the T cells. The higher affinity C24 T cells express *Ifng* and Il21 and skew towards a Th1 phenotype. The lower affinity C7 T cells skew towards a Th17 phenotype and are more dysfunctional. However, another factor to consider is that higher affinity T cells can outcompete the lower affinity T cells for antigen, raising the possibility that antigen availability in this model can modulate T cell phenotype. Finding that the two clones clustered separately, independently of TCR gene expression, demonstrates that TCR affinity influences T cell differentiation and function during chronic Mtb infection.

## Discussion

Following low-dose aerosol infection of B6 mice with virulent Mtb, the bacilli replicate for two to three weeks until T cells are recruited to the lung, which abrogates Mtb exponential growth and initiates the plateau phase of infection ^55^. In B6 mice, which are resistant to Mtb, the plateau phase can last for up to a year, during which time the mice appear healthy. During the terminal phase of infection, B6 mice lose weight for two to three weeks, and then expire. Why, after maintaining control for so long, the mice succumb to infection is unknown. We hypothesized that CD4 T cell dysfunction in chronically infected B6 mice contributes to loss of control of Mtb infection. We previously showed that T cell exhaustion occurs in susceptible C3H mice during Mtb infection ^56^. A confounder in characterizing T cell dysfunction is enumerating antigen-specific T cells. Generally, Mtb-specific T cells are counted only if they produce cytokines after antigen stimulation. Antigen stimulation induces TCR downregulation, which makes tetramer staining unreliable. Consequently, it difficult to ascertain whether non-cytokine producing T cells lack specificity for the antigen being tested, or whether they are dysfunctional. We call this the denominator problem, as the number of T cells that can potentially respond to antigen is unknown. Studies of CD8 T cell exhaustion during chronic viral infection or cancer solve this problem by using TCRtg or purified tetramer^+^ CD8 T cells ^13, 57, 58^. To study CD4 T cell dysfunction during chronic Mtb infection, we established an adoptive transfer model.

Transfer of C7 T cells into TCRα KO mice initially controls Mtb growth, and C7 T cells differentiate into polyfunctional effectors. By 25 wpi, bacterial recrudescence occurs, and the mice die early compared to B6 mice. At this time, C7 cytokine production diminishes compared to 4 wpi, and two T cell populations become dominant: one expressing high PD-1 (PD-1^hi^TIM-3^-^) levels and the other expressing PD-1 and TIM-3 (PD-1^hi^TIM-3^+^), resembling CD8 T_PEX_ and T_TEX_ cells, respectively ^11, 13^. These C7 T cell populations express gene signatures associated with T cell dysfunction and temporally associated with recrudescence. Representative genes include inhibitory receptors (*Lag3, Pdcd1, Tigit, Ctla4*), inhibitory cytokines (*Il10*), and *Hif1a*, which are expressed by exhausted CD4 T cells during chronic viral infection ^23, 59^. While the overall loss of IL-2, TNF, and IFNγ is consistent with exhaustion, high levels of *Ifng* mRNA persists in these cells, particularly in PD-1^hi^TIM-3^+^ cells. This functional state has been described for exhausted CD8 T cells during cancer and chronic viral infections. Given the importance of IFNγ in Mtb containment, it seems incongruous to label these T cells as dysfunctional or exhausted, although terminally exhausted CD8 T cells maintain their production of IFNγ^13^. However, IFNγ production correlates with bacterial load and too much IFNγ can exacerbate lung disease ^60^. Thus, IFNγ production by terminally exhausted CD4 T cells could be symptomatic of an inability to suppress Mtb replication. Instead of being beneficial, chronic IFNγ production could drive tissue inflammation. While diminished Th1 cytokine production could represent loss of function, it could also be a consequence of skewing towards a TFH-like phenotype (detected as increased *Il21* expression) as reported during chronic viral infections ^61^.

In B6 mice with an intact immune system, parenchymal CD4 T cells have increased coinhibitory receptor expression, reduced cytokine production and polyfunctionality, and decreased proliferation during chronic infection (i.e., > 40 wpi), all features of CD8 T cell exhaustion. Given the heterogeneity of T cell responses, we used scRNAseq to systematically analyze CD4 T cell function. After HN878 infection, Akter et al described four CD4 T cell clusters (naïve, Type I IFN responders, activated 1, activated 2) at 7-15 wpi ^62^. Our data, generated with a different Mtb strain and time points, also revealed clusters of cells with a naïve and type I IFN signature, as well as 11 additional CD4 T cell clusters. Remarkably, even 6 wpi, polyfunctional CD4 T cells were only a small fraction of the response. Four of the largest clusters had a core Th1 signature and extensively shared TCR clonotypes, indicating a common antigen specificity. TCR sharing was most extreme in Cluster 5: half of the TCR clonotypes in Cluster 5 were found in Cluster 2, indicating that the dominant clonotypes in these two distinct clusters share antigen specificity. While Cluster 5 CD4 T cells were polyfunctional, Cluster 2 CD4 T cells expressed lower levels of genes associated with anti-mycobacterial function (i.e., *Ifng, Tnf, Il21,* and *Tnfsf8*) and higher levels of co-inhibitory genes (i.e., *Entpd1, Tigit, Lag3,* and *Havcr2*). We infer that the recognition of Mtb-infected macrophages by Cluster 5 cells leads to their activation but ultimately these T cells become dysfunctional (i.e., exhausted) and reprogramming of their transcriptome leads to a distinct population of cells (i.e., Cluster 2). Particularly striking are the changes that occur in Clusters 2 and 5 between 6 wpi and 41 wpi. At 6 wpi, Both Clusters 2 and 5 are polyfunctional and express *Ifng, Tnf*, and *Il21*. By 41 wpi, only Cluster 2 CD4 T cells have largely loss *Tnf* and *Il21* expression. Relative to Cluster 2 and 5, Clusters 0 and 1 are hypofunctional with respect to cytokine production, which could arise from lack of antigen recognition ^63–66^ or inhibition by an inhibitory cytokine ^67–69^. TGFβ, which is made by macrophages in the lung^68, 70^ and by Tregs (Cluster 7), could suppress the function of cells in Cluster 0, 1, and 2, which express TGFβR mRNA. Interestingly, TGFβ signaling appears to be most relevant early during infection (6 wpi) and *Tgfbr2* expression is significantly reduced at 41 wpi. While the origin of the observed hypofunction in Cluster 0 and 1 is unknown, we speculate that it represents an inadequate response to persistent Mtb infection.

Diminished T cell function is caused by exhaustion (coinhibitory signals), anergy (deficient costimulation), senescence (blocked cell division), and inhibitory signals (IL-10, TGFβ). During chronic Mtb infection, pathways of UPR, ROS, and UV responses are upregulated, and these pathways can cause, or be symptomatic of, cellular stress, which also leads to T cell dysfunction ^71, 72^. Pathways associated with hypoxia, type I IFN signaling, and senescence are enriched in both PD-1^hi^TIM-3^-^ and PD-1^hi^TIM-3^+^ C7 cells. Type I IFN is associated with active TB in humans and is detrimental in murine TB ^73, 74^. A type I IFN signature raises the possibility that it contributes to T cell exhaustion, recrudescence, and disease development ^75–77^. In chronically infected mice, *Hif1a* is upregulated in C7 cells. Hif-1α is a critical regulator of cellular responses to changes in oxygen levels. Although Hif-1α is induced following TCR signaling, its upregulation can indicate that C7 cells are experiencing hypoxia ^78, 79^. Hypoxia is relevant to immune function as human TB granulomas become severely hypoxic ^80, 81^. Clusters 2 and 5 also include the cells most highly associated with a hypoxia gene signature, raising the possibility that the co-inhibitory receptors are being upregulated due to both metabolic stress and antigen stimulation. *Hif1a* expression and a hypoxia signature in association with impending recrudescence could indicate that metabolic stress affects CD4 T cell function. Indeed, the combination of hypoxia and chronic TCR stimulation synergistically causes CD8 T cell exhaustion in a tumor model ^78^. Type I IFN and metabolic stress induce cellular senescence, a state of cell cycle arrest caused by the cell cycle regulators p16, p21, and p53 ^16^. Originally described in fibroblasts ^82^, senescence also occurs in human T cells and results in altered function associated with susceptibility to viral infections and tumors ^16^. Senescent cells that develop a senescence-associated secretory phenotype (SASP) produce pro-inflammatory cytokines (IL-2, IL-6, IL-8, IFNγ, and TNF) and anti-inflammatory cytokines (IL-10 and TGF-β) ^16^. Senescence signatures were significantly enriched in PD-1^hi^TIM-3^-^ C7 cells, which could explain their reduced proliferation. Senescence should be considered as an alternate cause of T cell dysfunction during chronic TB.

In considering the factors that might affect T cell function, we investigated TCR affinity. After co-transfer of C7 and C24 cells, the higher affinity C24 cells resembled CD8 T_TEX_, with a PD-1^+^TIM-3^+^ phenotype and maintained the potential to produce IFNγ. In contrast, the lower affinity C7 cells resembled T_PEX_ (TIM-3^-^TCF-1^+^), and fewer cells could make IFNγ after stimulation. Interestingly, C24 and C7 cells both acquired a Th1 and Th17 signature, respectively. The divergence of phenotypes between the two clones was unexpected as individual T cell clones have the potential to differentiate into multiple effector cell types ^46^. We infer that higher affinity T cells might “protect” lower affinity clones from undergoing terminal differentiation and becoming terminally exhausted. The heterogeneity of the lung environment (i.e. antigen abundance) coupled with the polyclonal nature of T cell responses (i.e., TCR affinity) affects T cell differentiation and lead to varied T cell responses.

Our study provides crucial insights into the activation states of lung parenchymal CD4 T cells during TB and yields important findings that advance our understanding of CD4 T cell function and dysfunction during chronic bacterial infection. These include: (1) the loss of CD4 polyfunctionality during chronic Mtb infection is associated with higher lung CFU; (2) CD4 T cells that phenotypically resemble exhausted CD8 T cells are detected as early as 6 wpi and their increased frequency later during infection correlates with increased lung CFU; (3) cellular stress including hypoxia, type I IFN signaling, and senescence likely contribute to CD4 T cell dysfunction as manifested by senescence and exhaustion; and (4) polyclonal T cell responses appear to prevent the development of CD4 T cell dysfunction. There are several caveats to consider. First, the C7➔TCRα model is a reductionist approach and not all findings from this model were observed in C57BL/6 mice. Second, following aerosol Mtb infection, there is substantial variability in the onset of morbidity. Late time points were chosen based on when similar cohorts began to manifest disease. However, the mice evaluated in these studies were not obviously ill for logistical reasons, leading to uncertainty in the “clinical state” of the individuals analyzed. Third, we emphasized the parallels between dysfunctional CD4 T cells and exhausted CD8 T cells as there were striking similarities. However, the profile of dysfunctional Mtb-specific CD4 T cells does not fully align with standard definitions of exhausted CD8 T cells during cancer and chronic viral infection. The differences which we observe are potentially important as the definition of CD4 T cell exhaustion is still in flux ^23, 83, 84^. Finally, it is unknown whether CD4 T cell exhaustion impairs Mtb control. We previously showed that TIM-3 blockade enhances Mtb control ^56^; however, whether this is mediated by CD4 T cells remains to be established. Conversely, genetic deletion of PD-1 or PD-L1 leads premature death in mice, driven largely by pro-inflammatory CD4 T cells ^85^. Similarly, anti-PD-1 mAb treatment of rhesus macaques exacerbates TB ^86^. These studies suggest that PD-1 acts as a rheostat^87^ that modulates T cell responses and prevents tissue destruction^60^.

We propose a model in which increasing bacterial numbers lead to more antigen production and further TCR signaling. Together with cell extrinsic factors (e.g., hypoxia, type I IFN), chronic T cell activation causes T cell dysfunction and diminished ability to contain the infection. This permits more bacterial replication, and the cycle is repeated. While we are unable to determine the direction of causality, this feed-forward loop would result in recrudescence and disease. Alternatively, if we assume that exhaustion is a protective mechanism against the development of immunopathology, increased TCR signaling could overwhelm the ability of the PD-1 to moderate T cell activation, leading to increased inflammation. If, as observed during PD-1 blockade this results in increased Mtb burden^85, 86^, this could also result in a feed-forward loop. Further study is needed to ascertain whether reversing the effects of senescence or metabolic stress could protect immune responses without the potential complications of checkpoint blockade ^86, 88^.

## Acknowledgements

This work was funded by NIH/NIAID grants R01AI106725 and R01AI172905 to S.M.B; and NIH S10OD028576 for the purchase of the BD FACSFusion Cell Sorter. We thank the UMass Chan Flow Cytometry Core and Shayla Boyce for their expertise and excellent technical assistance. We acknowledge the NIH Tetramer Core Facility (contract number 75N93020D00005) for providing the ESAT6 tetramers.

## Author contribution

Conceptualization, E.C. and S.M.B.; Investigation, E.C., and K.C.; Formal analysis, E.C. and S.M.B.; Writing & Editing, E.C. and S.M.B.; Supervision, S.M.B.; Funding Acquisition, S.M.B.

## Supplemental Figures and Tables

**Table S1. Significant DEGs between Group B and Group C vs Group A C7 T cells. Tab 1**: Differentially expressed genes between Group A (uninfected; PD1^-^TIM3^-^) and Group B (4wpi; PD1^-^TIM3^-^) C7 cells. Criteria are log2FC>1 and padj<0.05. **Tab 2**: Differentially expressed genes between Group A (uninfected; PD1^-^TIM3^-^) and Group C (4wpi; PD1^+^TIM3^-^) C7 cells. Criteria are log2FC>1 and padj<0.05. **Tab 3**: GSEA output for Hallmark pathways enriched in Group A (uninfected; PD1^-^TIM3^-^) vs. Group B (4wpi; PD1^-^TIM3^-^) C7 cells. **Tab 4**: GSEA output for Hallmark pathways enriched in Group A (uninfected; PD1^-^TIM3^-^) vs. Group C (4wpi; PD1^+^TIM3^-^) C7 cells. **Tab 5**: Genes differentially expressed between Group A vs. Group B or C. Genes are special relevance are highlighted yellow.

**Table S2. Significant DEGs between Group B and Group C C7 T cells.** Differentially expressed genes between Group B (4wpi; PD1^-^TIM3^-^) and Group C (4wpi; PD1^+^TIM3^-^) C7 cells. Criteria are log2FC>1 and padj<0.05. Genes expressed more in Group C have a positive log2FC and are shaded beige. Genes expressed more in Group B have a negative log2FC and are shaded blue. Genes are special relevance are highlighted yellow.

**Table S3. GSEA Hallmark pathway enrichment results between Group C and Group D C7 T cells.** GSEA output for Hallmark pathways enriched in Group C (4wpi; PD1^+^TIM3^-^) vs. Group D (25wpi; PD1^+^TIM3^-^) C7 cells.

**Table S4. GSEA Hallmark pathway enrichment results between Group C and Group E C7 T cells.** GSEA output for Hallmark pathways enriched in Group C (4wpi; PD1^+^TIM3^-^) vs. Group E (25wpi; PD1^+^TIM3^+^) C7 cells.

**Table S5. Significant DEGs between Group C and Group D C7 T cells.** Differentially expressed genes between Group C (4wpi; PD1^+^TIM3^-^) and Group D (25wpi; PD1^+^TIM3^-^) C7 cells. Criteria are log2FC>1 and padj<0.05. Genes expressed more in Group D have a positive log2FC and are shaded beige. Genes expressed more in Group C have a negative log2FC and are shaded blue. Genes are special relevance are highlighted yellow.

**Table S6. Significant DEGs between Group C and Group E C7 T cells.** Differentially expressed genes between Group C (4wpi; PD1^+^TIM3^-^) and Group E (25wpi; PD1^+^TIM3^+^) C7 cells. Criteria are log2FC>1 and padj<0.05. Genes expressed more in Group E have a positive log2FC and are shaded beige. Genes expressed more in Group C have a negative log2FC and are shaded blue. Genes are special relevance are highlighted yellow.

**Table S7. Significant DEGs between Group D and Group E C7 T cells.** Differentially expressed genes between Group D (25wpi; PD1^+^TIM3^-^) and Group E (25wpi; PD1^+^TIM3^+^) C7 cells. Criteria are log2FC>1 and padj<0.05. Genes expressed more in Group E have a positive log2FC and are shaded beige. Genes expressed more in Group D have a negative log2FC and are shaded blue. Genes are special relevance are highlighted yellow.

**Table S8. GSEA Senescence signatures (related to Figure 2).** GSEA senescence signatures with Normalized Enrichment Score (NES) >|1.4| and a False Discovery Rate (FDR) <0.05 enriched in RNAseq data sets from Group C (4wpi; PD1^+^TIM3^-^) compared to Group D (25wpi; PD1^+^TIM3^-^) or Group E (25wpi; PD1^+^TIM3^+^) C7 cells.

**Table S9 (related to Figure 5). Differentially Expressed Genes (DEG) and cluster assignment for scRNAseq. Tab 1:** All Sig DEGs. All DEGs that have an adjusted p-value of <0.05. **Tab 2:** Top 30 Genes by Cluster. Top 30 DEGs in each cluster. Some of the DEGs in Clusters 6 and 12 do not have a padj<0.05. Clusters are color-coded to match Figure 5. Genes we consider immunologically significant are highlighted in yellow. **Tab 3:** DEG considered in Cluster assignment. **Tabs 4 – 16.** Each of these tabs lists the DEG for Clusters 0 – 12. All tabs have the same format. Column A–G list the DEG for the designated cluster compared to all other clusters and are from Tab 1 (all DEG have an adjusted p-value of <0.05). Column A, p_val; Column B, avg_log_2_FC; Column C, pct.1; Column D, pct.2; Column E, p_val_adj; Column F, cluster; Column G, gene. Some genes of immunological interest are highlighted beige (upregulated) or blue (downregulated). Column J–O list the DEG for the cluster when compared to Cluster 4 (naïve). These DEG have an adjusted p-value of <0.05, and are sorted by largest log_2_FC. **Tab 17:** a core signature expressed by Clusters 0, 1, 2, 3, 5, 6, and 9 compared to Cluster 4 (naïve T cells).

**Table S10 (related to Figure 6).** The results of Seurat’s FindMarker analysis to identify DEG for each cluster between 6 wpi vs. 41 wpi. The first tab is the analysis for all cells. Subsequent tabs are the analysis for each cluster. The data has been filtered for an absolute fold-change of >0.5 and an adjusted p-value of <0.05. A positive fold-change represents enrichment at 6 wpi and is displayed in a blue font. A negative fold-change represents enrichment at 41 wpi and is displayed in a red font.

**Table S11 (related to Figure 6). Panther pathway analysis.** Statistically significant DEGs that were upregulated at 6 or 41 wpi were found using Seurat FindMarkers (Table S10). These genes were used as input for Panther (pantherdb.org), which was used to determine molecular pathways that were statistically over-represented in the input list compared with randomly selected genes^42^. Tab 1: The gene input from Table S10 that was used as input. Subsequent tabs show the results for the Panther Overrepresentation analysis. All tabs are organized similarly. The first table identifies pathways in the Panther pathway collection. The second table, below, identifies pathways in the Reactome database.

**Table S12 (related to Figure 7). TCR data**. Original TCR data and analyses used in the paper (Excel spreadsheet). **Tab 1**: CD4_TCRs_all. Original TCR data displayed as a pivot table, **Tab 2**: analysis. Analysis of pivot table, focusing on the number of TCRs in each cluster, grouped by individual mouse, by time point, or by TCR expansion (i.e., clonotype). **Tab 3**: top 20 clonotypes AA. Clonal TCR expansions group by size and evaluated by cluster and mouse. Tope 20 clonotypes in each mouse individually. **Tab 4:** C7 CASSYWGGGQNTLYF. C7 CD4 T cells from uninfected transgenic mice were adoptively transferred to the B6 mice before infection and subsequent analysis. These ESAT6-specific CD4 T cells were identified based on their CDR3β nucleotide sequence which encodes the amino acids CASSYWGGGQNTLYF. All CD4 T cells with a CDR3β sequence of CASSYWGGGQNTLYF are listed. Subsequent analysis shows the associated CDR3α sequences, the distribution among different clusters, the clonal nature of the CDR3β nucleotide rearrangement, and the different CDR3α rearrangements.

**Table S13 (related to Figure 8). Differentially Expressed Genes (DEG) and cluster assignment for C7/C24 scRNAseq analysis.** Top 30 DEGs in each cluster. Clusters are color-coded to match Figure 1. Genes we consider immunologically significant for cluster assignment are highlighted yellow and other important genes are highlighted green. Some of the DEGs do not have a padj<0.05 (shaded pink).

## Methods

### Ethics statement

Studies were conducted using the relevant guidelines and regulations and approved by the Institutional Animal Care and Use Committee at the University of Massachusetts Medical School (UMMS) (Animal Welfare A3306-01), using the recommendations from the Guide for the Care and Use of Laboratory Animals of the National Institutes of Health and the Office of Laboratory Animal Welfare.

### Data Availability statement

All data supporting the findings of this study are available within the paper and its supplementary information. RNAseq data has been deposited to GEO. Further information and requests for resources and reagents should be directed to and will be fulfilled by the Lead Contact, Samuel Behar. Source data are provided with this paper.

### Mice

P25 TCRtg and TCRα KO mice were purchased from Jackson Lab (Bar Harbor, ME) and bred locally. C57BL/6J mice were purchased from Jackson Lab. C7 C57BL/6 (Vβ10; Thy1.1) and C24 RAG2^-/–^ (Vβ4; Thy1.2) TCRtg mice were originally obtained from Eric Pamer and bred locally^49, 90^. Experimental mice were 6 – 12 weeks old and sex-matched. For adoptive transfers sex-matched TCRα KO at 7-12 weeks old from different litters were randomly assigned to receive C7, C24, or both cells. All studies involving animals were conducted following relevant guidelines and regulations, and the studies were approved by the Institutional Animal Care and Use Committee at the University of Massachusetts Medical School (Animal Welfare A3306-01), using the recommendations from the Guide for the Care and Use of Laboratory Animals of the National Institutes of Health and the Office of Laboratory Animal Welfare.

### In vivo infections

The Erdman strain of *Mycobacterium tuberculosis*, which has been passaged through mice, was used for aerosol infection as previously described ^91^. Mice were infected through the aerosol route at a dose of ∼100 CFU/mouse using a Glas-Col chamber (Terre Haute, IN) as previously described ^18^. Frozen bacterial stocks were thawed, diluted into 5 mL of 0.01% Tween-80 in PBS, and sonicated for 1 minute prior to aerosolization. The average dose delivered into the lung was determined for each infection by plating lung homogenate from 4-5 mice 24 hours after infection and ranged between 40 – 150 CFU/mouse. Lung and spleen bacterial burden was determined using the left lung or whole spleen, which were harvested into 0.01% Tween-80 in PBS with 2 mm zirconium oxide beads (Next Advance). Tissue was homogenized using a FastPrep homogenizer (MP Biomedicals), serially diluted, and plated on 7H11 agar plates (Hardy Diagnosis). CFU was counted after 19-21 days. Morbidity studies were performed following infection. All mice were monitored weekly in accordance with IACUC guidelines using the Body Condition Score (BCS) and serial weight determinations. Mice with a BCS score less than or equal to 2 or had lost 20% of their maximum body weight were euthanized.

### TCRtg T cell Adoptive Transfer

T cells were isolated from the lymph nodes and spleens of C7 or C24 TCR transgenic mice. Single cell suspensions were made by passing spleens and lymph nodes through 70 µm and 40 µm filters sequentially, and a MojoSort^TM^ CD4 T cell isolation kit and magnet (Biolegend, San Diego, CA) was used. Cells were resuspended in 2% FBS in PBS, and 1 x 10^6^ cells/mouse were injected through the tail vein 24 hours before aerosol infection. Cell purities were determined for each experiment using flow cytometry.

### Intravascular Staining

1.2 - 2.4 µg/mouse fluorochrome-conjugated anti-CD45 antibody in 200 µL of 2% FBS in PBS was injected through the tail vein 2 minutes before euthanizing with CO_2_, and organs were harvested 1 minute later. Lungs were perfused and removed. Lymphocytes from blood collected in RPMI containing heparin (40 U/mL) were isolated using Lympholyte® (CEDARLANE) and analyzed by flow cytometry to confirm uniform staining.

### Cell Preparation

Lungs were perfused with 10 mL RPMI (Gibco) and collected into 5 mL complete RPMI with pen/strep (cRPMI + P/S; 10% FBS, 2 mM L-Glutamine, 100 units/mL Penicillin/Streptomycin, 1 mM Na-Pyruvate, 1X non-essential amino acids, 0.5X minimal essential amino acids, 25 mM HEPES, and 7.5 mM NaOH). Single cell suspensions were prepared by homogenizing lungs using a GentleMACS tissue dissociator (Miltenyi), digesting with collagenase (Sigma; 300 U/mL in cRPMI + P/S) at 37°C for 30 minutes, followed by a second run using the GentleMACS. Cell suspensions were then sequentially filtered through 70 µm and 40 µm strainers. Spleens were collected into cRPMI + P/S. Tissue was mechanically dissociated through 70 µm filter and washed with RPMI. Cell pellet was resuspended in ACK Lysis buffer and incubated at room temp for 1-2 minutes. After lysis, cells were washed with RPMI, and resulting cell pellet was resuspended in cRPMI and passed through a 40 µm filter to make a single cell suspension.

### Intracellular cytokine staining (ICS) stimulation and flow cytometric analysis

After processing, cells in single cell suspension were plated for stimulation or staining. Stimulation for ICS was done with anti-CD3 and anti-CD28 (Biolegend; each at 1 µg/mL) or ESAT-6 peptide (10 µM) for a total of 5 hours. See Table S14 for a complete list of antibodies and fluorochromes used in this study. Cells were stained with Zombie Fixable Viability dye (Biolegend) for 10 minutes at room temperature (RT). Some experiments used I-A^b^/ESAT-6_4-17_ and H2K^b^/Tb10.4_4-11_ MHC Tetramers alongside other antibodies. Class II tetramer staining was done at 37C for 1 hour, and surface and class I tetramer staining was done at 4C for 20 minutes. BD Cytofix/Cytoperm Kit was used for ICS. Transcription factor (TF) staining was done with eBiosciences FoxP3 Transcription Factor staining kit, where cells were fixed and permeabilized for 30 minutes at RT, followed by staining with antibodies against various TFs for another 30 minutes at RT. Samples were fixed for a total of 1 hr with 1% - 4% PFA, washed, and resuspended in autoMACS running buffer and were acquired on Aurora (Cytek) or MACSQuant (Miltenyi) flow cytometers. Data was analyzed using FlowJo v10.8.1. and SPICE v6.1 ^92^. I- A^b^/ESAT-6_4-17_ and H2K^b^/Tb10.4_4-11_ MHC Tetramers were produced by the NIAID Tetramer Core (Emory, Atlanta, Georgia).

### Cell Sorting

Lung cells were prepared as described, with a red blood cell lysis step prior to passing cell suspension through 40 µm filter. Cell pellet was resuspended in 1 mL ACK Lysis Buffer (GIBCO) for 1-2 minutes at room temp. Cells were washed with RPMI then resuspended in media and passed through 40 µm filter to make single cell suspension. Cells were stained with flow antibodies prior to sorting on FACSAria II (BD Biosciences) or MA900 (Sony) cell sorters.

### RNAseq

Parenchymal lung C7 T-cells were sorted from the lungs of infected mice at 4 and 25 wpi. CD4 T cells were identified based on the following scheme: lymphocyte sized gate ➔ single cells ➔ dead and dump (NK1.1/CD19/TCRγδ) negative ➔ CD8^-^ ➔ CD45-IV^-^ ➔ CD4^+^Vβ10^+^. These cells were then sorted based on their expression of TIM-3 and PD-1 and placed in RNAprotect (QIAGEN). RNA was isolated using RNeasy kit (QIAGEN). Sequencing, alignment, and processing of data was performed by GENEWIZ. Differential expression analysis was performed in R (v4.3.0) using DESeq2 (v1.40.2). GSEA was performed using the Broad Institute software Gene enrichment scores were determined using the Gene Set Enrichment Analysis (GSEA) desktop app (Mac version 4.3) produced by the Broad Institute ^93, 94^. The mouse Hallmark database was primarily used ^95^. The RNAseq data has been deposited at GEO and are publicly available as of the date of publication (Accession number: GSE266006).

### Single-cell RNAseq Library Generation and Sequencing

Total parenchymal lung t-cells were sorted from the lungs of Mtb-infected B6 mice at 6 and 41 wpi (6 wpi; 1 x 10^5^ or 41 wpi; 1 x 10^6^ C7 cells IV transferred 16-24 hours before aerosol infection) and C7/C24 ➔ TCRα KO mice at 26 wpi. Sorted cells were processed and cDNA libraries were made according to 10x Genomics 5’ Single Cell Immune Profiling protocols. Cells were loaded in chips to capture 10,000 cells/lane, and the resulting GEM droplets underwent reverse transcription to label transcripts with unique molecular identifiers (UMIs). Samples were then removed from the BSL-3 space, and GEX and V(D)J libraries were performed using manufacturer’s instructions. 13 rounds of PCR cycles were performed for cDNA amplification, and 14 rounds of cycles were performed for GEX Sample Index PCR. Libraries were sequenced on an Illumina NextSeq 500, and 5,000 and 20,000 read pairs per cell were targeted for V(D)J and GEX libraries, respectively.

### Single-cell RNAseq and TCRseq Data Processing

Sequencing data was processed and aligned against the mouse reference genome mm10 using Cellranger v6 on the UMass High Performance Computing SCI Cluster. The resulting gene count matrices and filtered contig sequences were further analyzed using R (v4.3.0) and Seurat (v4.3.0). Cells with greater than 10% mitochondrial genes were filtered out. The TCR genes were removed from the gene expression data before dimensionality reduction and Louvain clustering to avoid TCR bias among clonally expanded T cells affecting the outcome. Samples were then normalized using default parameters, and most variable genes were detected using the FindVariableFeatures function. The data was then scaled using ScaleData, and principal component analysis (PCA) was done using RunPCA. Clustering was performed using FindNeighbors (PCs 1:20), FindClusters, and RunUMAP (PCs 1:20). DoubletFinder (v2.0.3) was then used to find and filter out doublets in the samples. CD4s were then subsetted from the dataset for further analysis (Cd8a < 1e-10 & Cd3e > 1 & Cd4 > 0.5 & Cd8b1 < 1e-10) for the B6 analysis. C7 and C24 t-cells were subsetted using the CDR3β sequence (C7: CASSYWGGGQNTLYF; C24: CASSRQGGNYAEQFF). Samples were integrated using SelectIntegrationFeatures, FindIntegrationAnchors, and IntegrateData, and TCR information was merged into the Seurat object using scRepertoire (v1.10.1). The cells in the integrated Seurat object were clustered using FindClusters (resolution 0.6), and FindAllMarkers was used to determine the top defining genes of each cluster (min.pct = 0.25, logfc.threshold = 0.25). One cluster in each experiment was enriched in lncRNAs (Gm26917 and Gm42418) associated with ribosomal RNA contamination and excluded from further analysis. Cells were re-clustered using FindClusters (resolution 0.6). For polyclonal cells, FindMarkers (default parameters) was used to compare gene expression of cells in selected clusters against the Naïve cluster. The function getCirclize from scRepertoire was used to visualize TCR overlap between clusters, and repOverlap from Immunarch (v0.9.0) was used to calculate the Jaccard Similarity Index for TCRs between each cluster pair. All other TCR analyses were done using Pivot tables in Excel. Several other R packages within Tidyverse and ggplot2 were used for intermediate data processing steps and data visualization. The scRNAseq data have been deposited at GEO and are publicly available as of the date of publication (Accession number: GSE265891).

### Statistical Analysis

Statistical analysis was performed using Prism 10 (GraphPad). P-values were calculated using unpaired two-sided t test, one-way ANOVA, or as indicated in the figure legends. For normally distributed data an ordinary one-way ANOVA was used followed by Bonferroni’s multiple comparisons test. For two-way ANOVA, Bonferroni, Sidak’s, Tukey’s or Fisher’s Least Significant Difference multiple comparison test was used.

---

**Figure S1.**
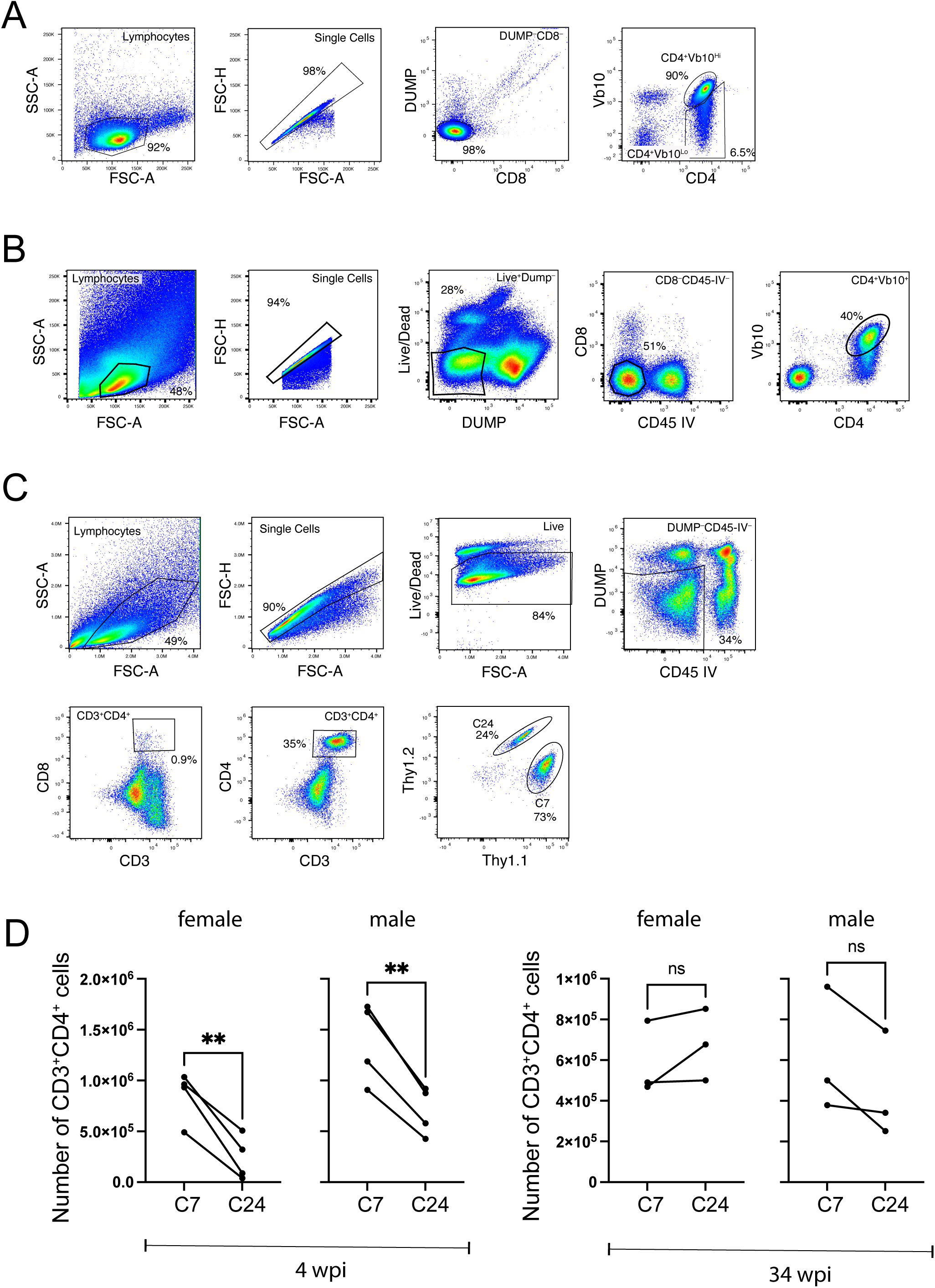
Gating strategy for flow cytometric analysis of C7 or C24 TCRtg CD4 T cells in the adoptive transfer model. (A) Analysis of CD4 T cells purified from C7 mice. Purity of CD4 T cells were between 90-95%. Between 70-95% of the CD4 T cells were identified as C7 TCRtg CD4 T cells based on staining with anti-Vβ10 or ESAT6 tetramers. (B) Gating scheme used to sort parenchymal (i.e., CD45-IV^-^) C7 TCRtg CD4 T cells from Mtb-infected mice after adoptive transfer. Sort gates are shown in Figure 3. (C) Gating scheme of flow cytometry analysis of the C7/C24 co-transfer experiments parenchymal. After gating on viable parenchymal (i.e., CD45- IV^-^) CD4 T cells, C7 and C24 TCRtg T cells were identified based on expression of Thy1.1 (C7) and Thy1.2 (C24). (D) The number of viable lung parenchymal Thy1.1 (C7) and Thy1.2 (C24) CD4 T cells in the lung at 4 and 34 weeks after adoptive transfer into TCRα knockout mice and Mtb infection. **, p < 0.01; ns, not significant by paired t-test.

**Figure S2.**
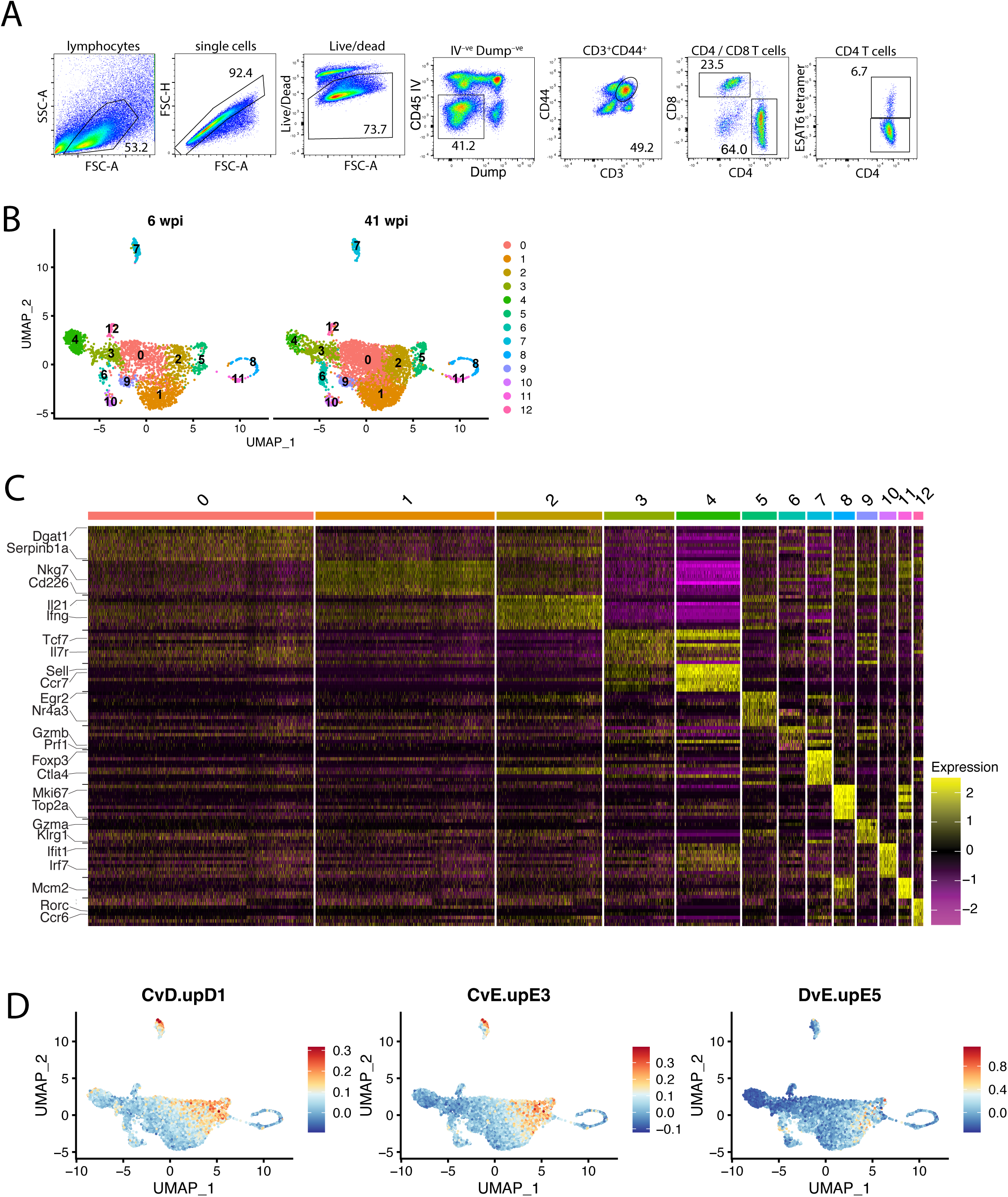
(related to Figure 5). Heat map of cluster-defining genes in B6 scRNAseq. (A) Gating scheme of flow cytometry analysis of parenchymal (i.e., CD45-IV^-^) ESAT6-specific CD4 T cell responses. (B) UMAP projection of CD4 T cells 6 and 41 wpi were analyzed by scRNAseq, integrated, and then split by timepoint. (C) Heatmap of top 10 genes within each cluster. Cluster-defining genes of interest are listed. (D) AddModuleScore plot using C7 RNAseq DEGs as gene signatures and applying to B6 scRNAseq dataset. Clusters 2 and 5 were most highly associated with the genes significantly upregulated in Group D and E compared to Group C. Cluster 2 expressed genes upregulated in Group E compared to Group D, which are associated with T cell exhaustion.

**Figure S3.**
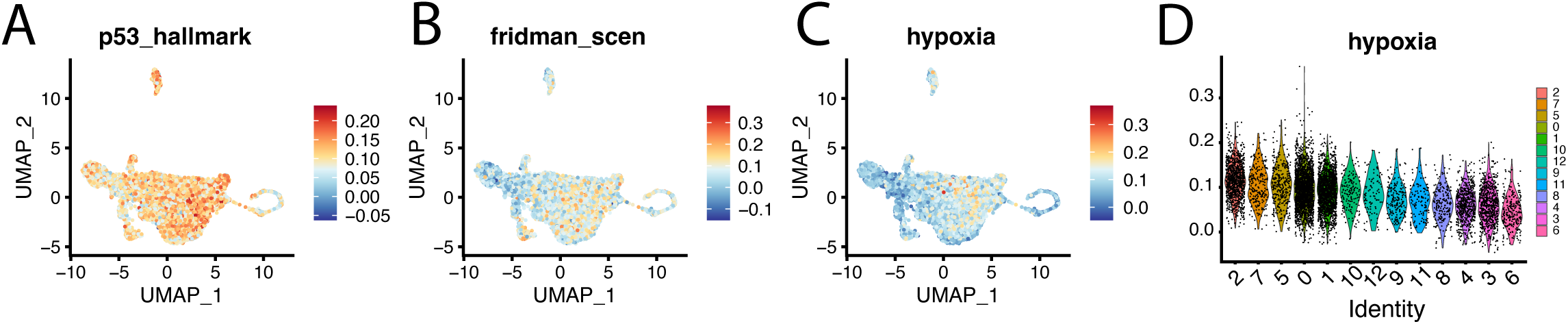
(related to Figure 5). Visualization of module scores calculated by Seurat’s *AddModuleScore* function using MSigDB gene sets. Feature plots (A-C) or violin plot (D) of the module scores calculated using the p53 Hallmark (A), Fridman Scen (B), and Hallmark Hypoxia (C-D) gene sets.

